# A skeletal muscle–sympathetic nerve–intestine network underlies muscle inflammation and atrophy induced by immobilization

**DOI:** 10.1101/2025.02.25.640219

**Authors:** Yu Hirata, Kazuhiro Nomura, Kaori Hozumi, Kenji Sugawara, Yusei Hosokawa, Kana Uchiyama, Tomoya Inoue, Tomoko Nishigaki, An Liyuan, Serika Yamada, Yutaro Miyatake, Takahiro Niikura, Tomoaki Fukui, Keisuke Oe, Ryosuke Kuroda, Sotaro Hamakita, Shohei Kumagai, Jun Hirata, Gerard Karsenty, Yushi Hirota, Kento Ohbayashi, Toshihiko Yada, Kazunari Miyamichi, Christopher K. Glass, Yusaku Iwasaki, Wataru Ogawa

## Abstract

Immobility is a common cause of muscle atrophy, but the underlying mechanisms have remained unclear. Here we show that limb immobilization in mice elicits inflammation and atrophy of skeletal muscle that are preventable by neutralizing antibodies to the chemokine CXCL10. Limb immobilization also induced changes to the gut microbiota and intestinal inflammation, and either sterilization of the intestine with antibiotics or administration of 10-hydroxy-*cis*-12-octadecenoic acid—a linoleic acid–derived gut microbial metabolite—prevented intestinal and muscle inflammation as well as muscle atrophy induced by immobilization, implicating intestinal inflammation in muscle inflammation and atrophy. Limb immobilization activated sympathetic nerves and increased β2-adrenergic receptor gene (*Adrb2*) expression in the intestine. Single-cell RNA-sequencing analysis revealed that, among cells expressing *Adrb2* in the intestine, immobilization increased only the population of macrophages. Pharmacological inhibition or macrophage-specific ablation of Adrb2 prevented immobilization-induced intestinal and muscle inflammation. Our results thus implicate a previously unrecognized muscle-nerve-intestine network in immobilization-induced muscle atrophy.

## Introduction

Muscle mass is an important determinant of human health. A reduction in muscle mass together with the consequent decline in physical performance can trigger or accelerate various pathological conditions.^1^ Exercise is an important stimulus for muscle growth,^2^ and immobilization is one of the most common causes of muscle mass loss.^3^ The decline in muscle mass induced by immobilization is attributable at least in part to the loss of exercise stimulation.^4^ However, whereas the exercise-induced increase in muscle mass generally occurs over a time span of weeks to months, immobilization elicits a more rapid decline in muscle mass, sometimes within just a few days,^5^ suggesting that immobilization actively triggers mechanisms that reduce muscle mass in a manner independent of the loss of exercise-induced muscle growth. The mechanisms by which immobilization proactively induces muscle mass loss have remained unclear, however. We recently showed that limb immobilization, either by cast-fixation or motor nerve resection, results in a decline in the basal intracellular Ca^2+^ concentration of muscle cells through downregulation of the Ca^2+^ channel Piezo1.^6^ This reduction in the cytosolic Ca^2+^ level in turn promotes protein catabolism in a manner dependent on the transcription factor KLF15 and the cytokine interleukin (IL)–6. These findings suggest that an increase in protein catabolism mediated by a Piezo1/KLF15/IL-6 pathway is a key contributor to the loss of muscle mass actively triggered by immobility.^6^ On the other hand, inflammatory signals have also been implicated in immobility-induced muscle atrophy,^7,8,9^ although the relation between protein catabolism and inflammation during immobility has remained unclear.

To provide further insight into the mechanisms by which immobility induces muscle atrophy, we have now investigated the time-dependent changes in skeletal muscle during limb immobilization. We found that the prominent pathological effect of limb immobilization on skeletal muscle transitions from protein catabolism to inflammation in a time-dependent manner. We also found that this muscle inflammation is dependent on changes to the gut microbiota and on intestinal inflammation triggered by the activation of sympathetic nerves in the intestine.

## Results

### Transition from protein catabolism to inflammation during skeletal muscle immobilization

We previously showed that limb immobilization for a relative short period (up to 3 days) induced by either cast-fixation or motor nerve resection results in a decline in skeletal muscle mass through an increase in protein catabolism mediated by a Piezo1/KLF15/IL-6 axis.^6^ We here investigated how more prolonged limb immobilization affects skeletal muscle. Cast-fixation of the hind limbs of mice resulted in a significant reduction in muscle mass of the immobilized limbs within the first 3 days of immobilization, and this decline in muscle mass continued thereafter in a time-dependent manner (Figure 1A). Prolonged immobilization did not affect food intake, body mass, or epididymal white adipose tissue (eWAT) mass (Figures S1A–S1C). The progression of muscle fiber atrophy during immobilization was confirmed by a decrease in muscle fiber area detected by histological analysis (Figures 1B and 1C, Figures S1D and S1E). The decrease in fiber area was observed in both slow- and fast-twitch fibers (Figures S1F and S1G). Immunohistochemical staining revealed that limb immobilization for 10 days, but not that for 3 days, resulted in a significant accumulation of cells positive for the macrophage marker F4/80 in the affected muscle (Figures 1B and 1D, Figures S1D and S1H). Flow cytometry also revealed an increase in the proportion of CD11b^+^F4/80^+^ cells among CD45^+^Ly6G^−^ cells in muscle (Figures 1E and 1F), and that the percentages of CD11c^+^CD206^−^ and CD11c^−^CD206^+^ cells among these CD11b^+^F4/80^+^ cells were increased and decreased, respectively, after limb immobilization for 10 days (Figures S2A–S2C). These results thus indicated that limb immobilization for 10 days induced the accumulation of macrophages in skeletal muscle as well as increased and decreased the proportions of pro-inflammatory and anti-inflammatory subsets of these cells, respectively.

**Figure 1.**
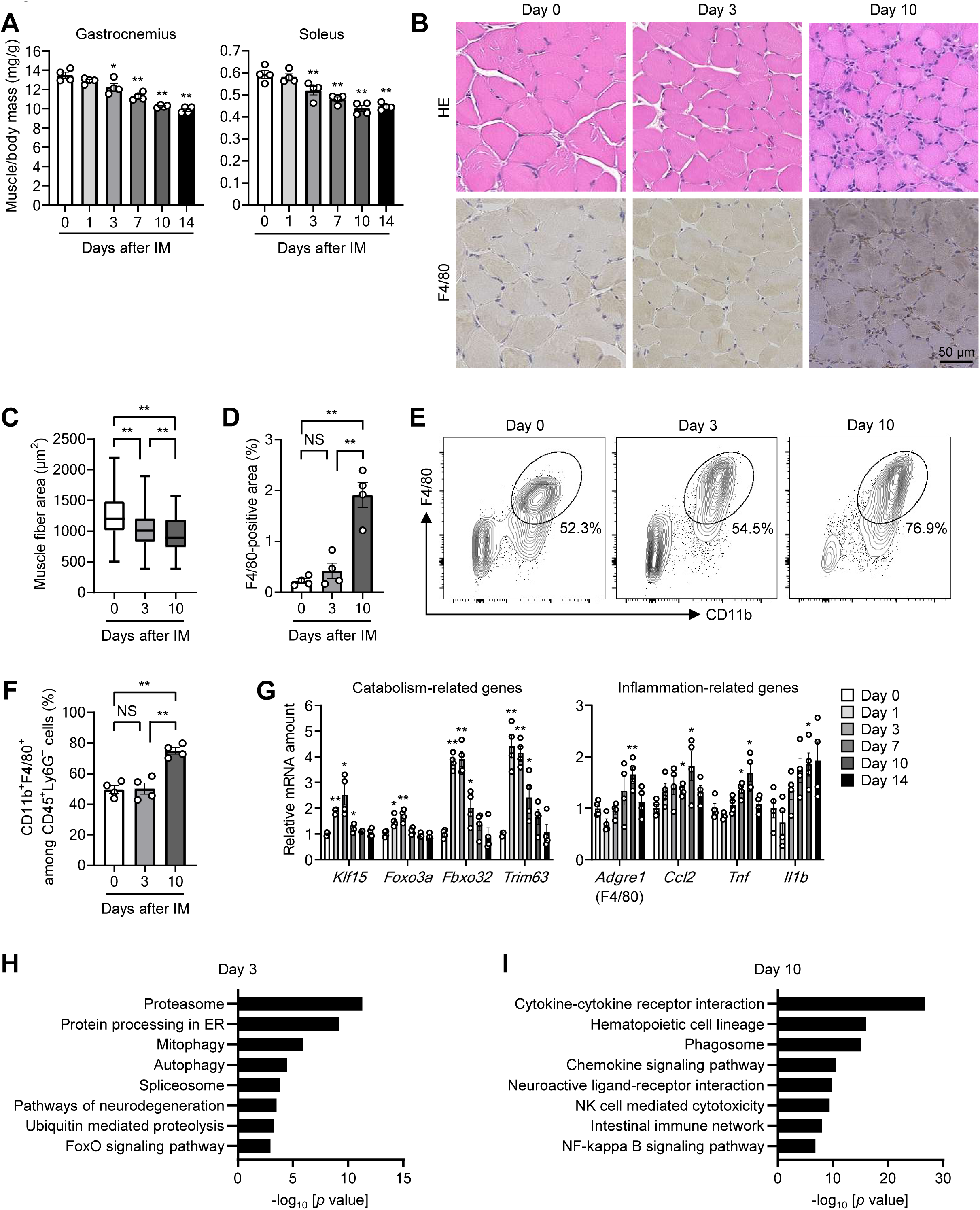
Transition from protein catabolism to inflammation during skeletal muscle immobilization. (A) Ratio of gastrocnemius and soleus muscle mass for both hind limbs to body mass for mice subjected to bilateral hind limb immobilization (IM) by cast-fixation for the indicated times (*n* = 4 mice). (B–D) Hematoxylin-eosin (HE) staining and immunohistochemical staining of F4/80 (B) for determination of muscle fiber area (C) and the F4/80-positive area (D) in soleus muscle of mice subjected to hind limb immobilization for the indicated times (*n* = 4 mice). The area of 800 fibers pooled from four mice was measured and averaged for each condition in (C). Scale bar (B), 50 μm. (E and F) Representative flow cytometric analysis (E) and the percentage (F) of CD11b^+^F4/80^+^ cells among CD45^+^Ly6G^−^ cells in immobilized skeletal muscle (*n* = 4 mice). (G) RT-qPCR analysis of protein catabolism– or inflammation-related gene expression in gastrocnemius muscle of immobilized mice (*n* = 4 mice). (H and I) Kyoto Encyclopedia of Genes and Genomes (KEGG) pathway analysis for RNA-seq data from the gastrocnemius at 3 days (H) or 10 days (I) after hind limb immobilization. The top eight pathways enriched among genes whose expression was upregulated by immobilization are shown. Quantitative data are presented as means ± SEM (A, D, F, and G) or medians with interquartile range (C). \**p* < 0.05, \*\**p* < 0.01, NS (not significant) compared with day 0 (A and G) or as indicated (C, D, and F) by two-way ANOVA with the Bonferroni post hoc test.

Reverse transcription and quantitative polymerase chain reaction (RT-qPCR) analysis revealed that the expression of genes related to protein catabolism was upregulated in immobilized muscle, peaking at 1 to 3 days (Figure 1G and Figure S2D), consistent with our previous findings.^6^ In contrast, the expression of genes related to inflammation was significantly upregulated during a later phase of immobilization, peaking at 10 days (Figure 1G and Figure S2D). RNA–sequencing (seq) analysis of immobilized muscle indicated that genes related to the proteasome, protein processing in the endoplasmic reticulum (ER), mitophagy, and autophagy were upregulated at 3 days (Figure 1H), whereas those related to cytokine–cytokine receptor interaction, hematopoietic cell lineage, phagosome, and chemokine signaling pathway were upregulated after immobilization for 10 days (Figure 1I). Gene set enrichment analysis (GSEA) also confirmed the upregulation of genes related to the proteasome or to cytokine–cytokine receptor interaction after 3 and 10 days of immobilization, respectively (Figures S2E and S2F).

Hind limb immobilization induced by motor nerve denervation similarly resulted in a time-dependent decline in skeletal muscle mass that was associated with the upregulation of genes related to protein catabolism or to inflammation at 3 and 10 days, respectively, as well as with the accumulation of F4/80-positive cells at 10 days (Figure S3). Together, these results thus revealed a transition from protein catabolism to inflammation during muscle atrophy triggered by prolonged limb immobilization.

### CXCL10 mediates inflammation and atrophy of muscle induced by immobilization

We next attempted to identify a humoral factor that might contribute to muscle inflammation during limb immobilization. RNA-seq analysis indicated that the gene for the chemokine CXCL10 was one of the most markedly induced inflammatory humoral factor genes in skeletal muscle under this condition (Figures 2A and 2B). The expression of *Cxcl10* was increased 1 day after the onset of limb immobilization by cast-fixation, peaked at 7 days, and decreased thereafter (Figure 2C). The circulating (Figure 2D) and muscle (Figure S4A) concentration of CXCL10 was similarly increased during limb immobilization. The upregulation of *Cxcl10* expression was apparent in both the blood cell and non–blood cell fractions of muscle, which were discriminated on the basis of the expression of CD45 (Figure S4B); however, it was more pronounced in the non–blood cell fraction, which was composed mostly of myofibers. These results, along with the finding that CXCL10 concentrations in muscle tissue were approximately ten times greater than those in circulation (Figure 2D and Figure S4A), suggest that the cytokine acts in an autocrine/paracrine manner. Moreover, a similar increase in *Cxcl10* expression was also detected in skeletal muscle of denervated mice (Figure S4C). Of note, upregulation of the CXCL10 gene (Figure 2E) and of other inflammation-related genes (Figure S4D) was observed in human skeletal muscle after cast-fixation for an average of 8.3 ± 3.1 days as a result of bone fractures. Furthermore, analysis of publicly available human gene expression data revealed that acute immobilization of lower limbs was associated with an increase in the abundance of *CXCL10* mRNA in skeletal muscle (Figure S4E).

**Figure 2.**
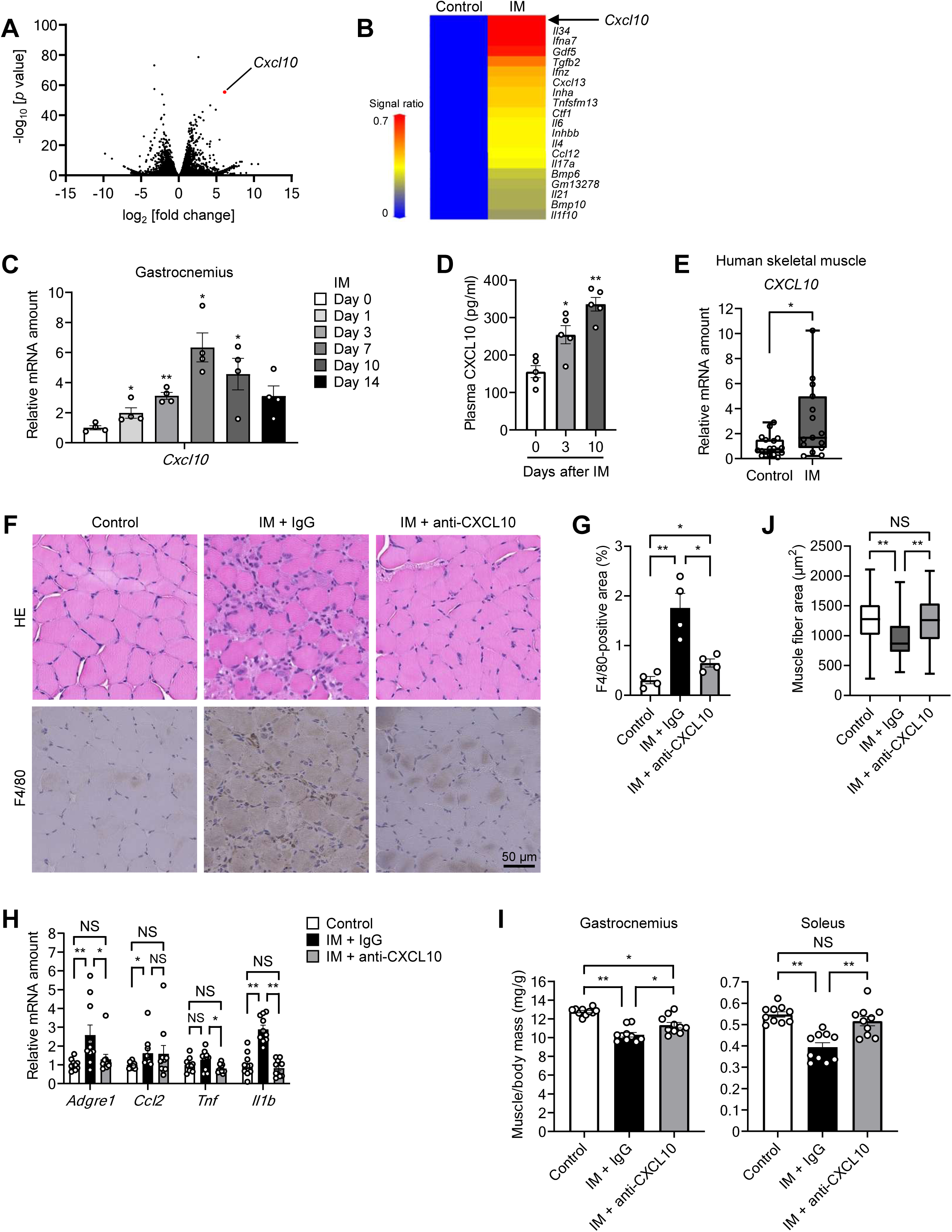
CXCL10 mediates inflammation and atrophy of muscle induced by limb immobilization. (A and B) Volcano plot of RNA-seq data (A) and heat map of inflammation-related gene expression determined by DNA microarray analysis (B) for gastrocnemius of mice subjected to cast-immobilization for 3 days compared with control mice. (C) RT-qPCR analysis of *Cxcl10* mRNA abundance in gastrocnemius of cast-immobilized mice (*n* = 4 mice). (D) Plasma concentration of CXCL10 in cast-immobilized mice (*n* = 5 mice). (E) RT-qPCR analysis of *CXCL10* mRNA in skeletal muscle of human individuals subjected to cast-fixation (*n* = 15) or of control subjects (*n* = 18). (F–J) Hematoxylin-eosin and F4/80 immunohistochemical staining (F) and the F4/80-positive area (G) of soleus (*n* = 4 mice), RT-qPCR analysis of inflammation-related gene expression in gastrocnemius (*n* = 10 mice) (H), the ratio of muscle mass to body mass (*n* = 10 mice) (I), and muscle fiber area in soleus (*n* = 4 mice) (J) for control mice or mice subjected to intraperitoneal injection of neutralizing antibodies to CXCL10 (0.1 mg/body) or control immunoglobulin G (IgG) 3 days before cast-immobilization for 10 days. Scale bar (F), 50 μm. The area of 800 fibers pooled from four mice was measured and averaged for each condition in (J). Quantitative data are presented as means ± SEM (C, D, G, H, and I) or medians with interquartile range (E and J). \**p* < 0.05, \*\**p* < 0.01, NS relative to day 0 (C and D) or as indicated (E and G–J) by the Mann-Whitney *U* test (E) or two-way ANOVA with the Bonferroni post hoc test (C, D, and G–J).

The administration of neutralizing antibodies to CXCL10 inhibited both the accumulation of macrophages (Figures 2F and 2G) and the upregulation of inflammation-related gene expression (Figure 2H) in muscle during subsequent limb immobilization by cast-fixation. These antibodies also attenuated the decrease in both skeletal muscle mass (Figure 2I) and muscle fiber area (Figures 2F and 2J) induced by prolonged limb immobilization. Immobilization increased the phosphorylation of ERK, a signaling molecule activated by CXCL10,^10^ in the gastrocnemius muscle, which was prevented by the neutralizing antibodies (Figure S5A). These results suggest that the antibodies indeed attenuated CXCL10-mediated signaling in muscle. The neutralizing antibodies to CXCL10 did not affect the upregulation of catabolism-related genes or the decline in muscle mass apparent after limb immobilization for 3 days (Figures S5B and S5C).

Collectively, these findings suggested that inflammation mediated by CXCL10 plays a key role in the loss of muscle mass induced by immobilization, whereas protein catabolism, which is activated relatively early during immobilization, is independent of this CXCL10 pathway.

### Prolonged limb immobilization induces inflammation in the intestine

Given that prolonged limb immobilization gives rise to inflammation in skeletal muscle, we next explored whether such immobilization also induces inflammation in other organs. Examination of the entire body of cast-immobilized mice revealed intestinal shortening (Figures 3A and 3B), a decrease in cecum mass (Figure 3C), and a reduction in the depth of crypts in the colon (Figures 3D and 3E), all of which are associated with intestinal inflammation.^11^ A decrease in tissue mass was also apparent for the small and large intestine, but not for the stomach or duodenum (Figure S6A). The reduction in the length of the small and large intestine as well as that in cecum mass became evident within 7 to 10 days of the onset of limb immobilization (Figures S6B and S6C), showing a time course similar to that of skeletal muscle inflammation.

**Figure 3.**
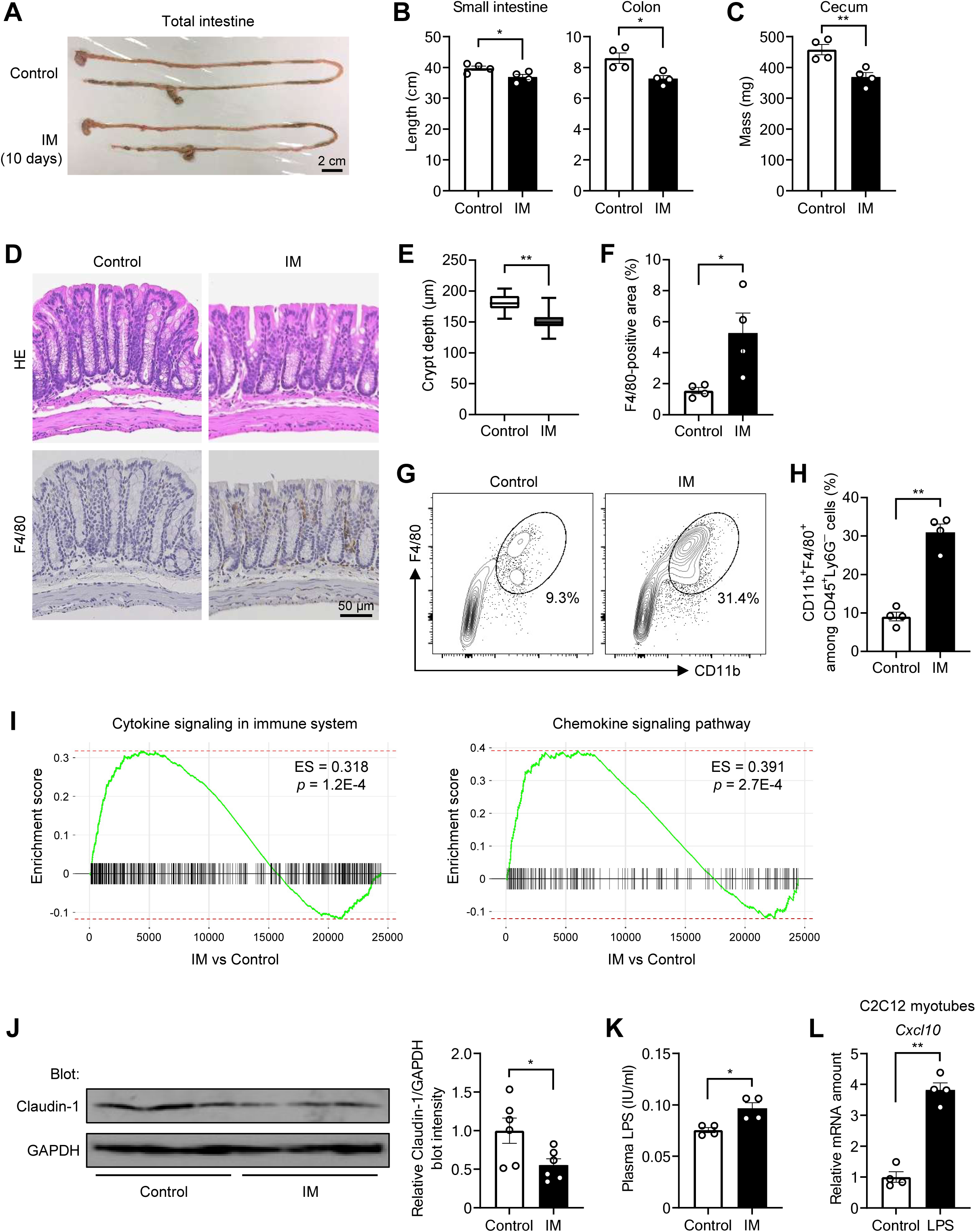
Prolonged limb immobilization induces inflammation in the intestine. (A–F) Representative images of the entire intestine (A), length of the small intestine and colon (B), mass of the cecum (C), and hematoxylin-eosin and F4/80 immunohistochemical staining (D) for determination of crypt depth (E) and the F4/80-positive area (F) in the colon of control mice or mice subjected to cast-immobilization for 10 days (*n* = 4 mice). Scale bars, 2 cm (A) and 50 µm (D). The depth of 100 crypts pooled from four mice was measured and averaged for each condition in (E). (G and H) Representative flow cytometric analysis (G) and the percentage (H) of CD11b^+^F4/80^+^ cells among CD45^+^Ly6G^−^ cells in the colon of immobilized and control mice (*n* = 4 mice). (I) GSEA of RNA-seq data obtained from the colon of immobilized and control mice. (J and K) Immunoblot analysis of claudin-1 and GAPDH (loading control) in the colon (*n* = 6 mice) (J) and the plasma concentration of LPS (*n* = 4 mice) (K) for mice subjected (or not) to cast-immobilization for 10 days. (L) RT-qPCR analysis of *Cxcl10* mRNA abundance in C2C12 myotubes exposed to LPS (1 μg/ml) or vehicle (control) for 6 h (*n* = 4 independent experiments). Quantitative data are presented as means ± SEM (B, C, F, H, and J–L) or medians with interquartile range (E). \**p* < 0.05, \*\**p* < 0.01 (unpaired Student’s *t* test).

Immunohistochemical analysis and flow cytometry revealed that prolonged limb immobilization resulted in the accumulation of F4/80-positive cells and CD11b^+^F4/80^+^ cells, respectively, in the intestinal wall (Figures 3D and F–H), with a concomitant increase in the proportion of pro-inflammatory (CD11c^+^CD206^−^) and decrease in that of anti-inflammatory (CD11c^−^ CD206^+^) macrophages (Figures S7A–S7C). In addition, RT-qPCR analysis showed that the expression of inflammation-related genes, was upregulated in the distal small intestine and colon, but not in the proximal small intestine, of mice subjected to cast-fixation for 10 days (Figure S8A). Cast-fixation did not upregulate *Cxcl10* in the small or large intestine (Figure S8B).

GSEA of RNA-seq data obtained from the intestine of such mice revealed upregulation of genes related to inflammation, including those associated with cytokine and chemokine signaling pathways (Figure 3I). Denervation-induced limb immobilization for 10 days also resulted in intestinal shortening (Figure S8C), a decrease in cecum mass (Figure S8D), and upregulation of inflammation-related genes in the distal small intestine and colon (Figure S8E).

Intestinal inflammation is typically associated with impairment of the intestinal barrier, resulting in an increase in the circulating levels of bacterial components such as lipopolysaccharide (LPS).^12^ Consistent with this scenario, prolonged limb immobilization resulted in a decrease in the intestinal abundance of claudin-1 (Figure 3J), a key protein in maintenance of the intestinal barrier,^13^ and in an increase in the plasma concentration of LPS (Figure 3K). Furthermore, treatment of cultured C2C12 myotubes with LPS resulted in the upregulation of *Cxcl10* expression (Figure 3L).

Collectively, these results suggested that prolonged limb immobilization triggers intestinal inflammation, which may then contribute to upregulation of CXCL10 expression and consequent induction of inflammation in skeletal muscle. Given the increase in plasma LPS concentration, we analyzed the effect of hindlimb immobilization on other skeletal muscles, including forelimb muscles (flexor carpi ulnaris) and diaphragm. Whereas the mRNA abundance of *Il1b* was increased (Figure S9A), the expression of *Cxcl10* (Figure S9A), as well as the mass (Figure S9B) of these muscles, remained unaltered, suggesting that inflammation occurred systemically, but another factor induced in immobilized muscles is required for the upregulation of *Cxcl10* and muscle atrophy. In addition, RNA-seq analysis of the forelimb muscles showed the upregulation of genes related to muscle contraction, exercise, and amino acid metabolism (Figure S9C), suggesting that hindlimb immobilization may induce a compensatory response in the forelimb muscles.

### The gut microbiota contributes to inflammation and atrophy of skeletal muscle induced by limb immobilization

The development of intestinal inflammation is closely related to changes in the gut microbiota.^12^ We therefore investigated the impact of prolonged limb immobilization on the gut microbiota. Analysis of bacterial 16S rRNA genes revealed that prolonged immobilization of the hind limbs of mice by cast-fixation resulted in marked changes to the gut microbiota including an increase in the abundance of the Lachnospiraceae group and a decrease in that of *Lactobacillus* species (Figures 4A and 4B). These changes in the gut microbiota were also demonstrated by principal coordinate analysis (Figure 4C), and they were largely reversed within 2 weeks after cast removal (Figures S10A–S10C), indicating that the changes were indeed primarily triggered by limb immobilization.

**Figure 4.**
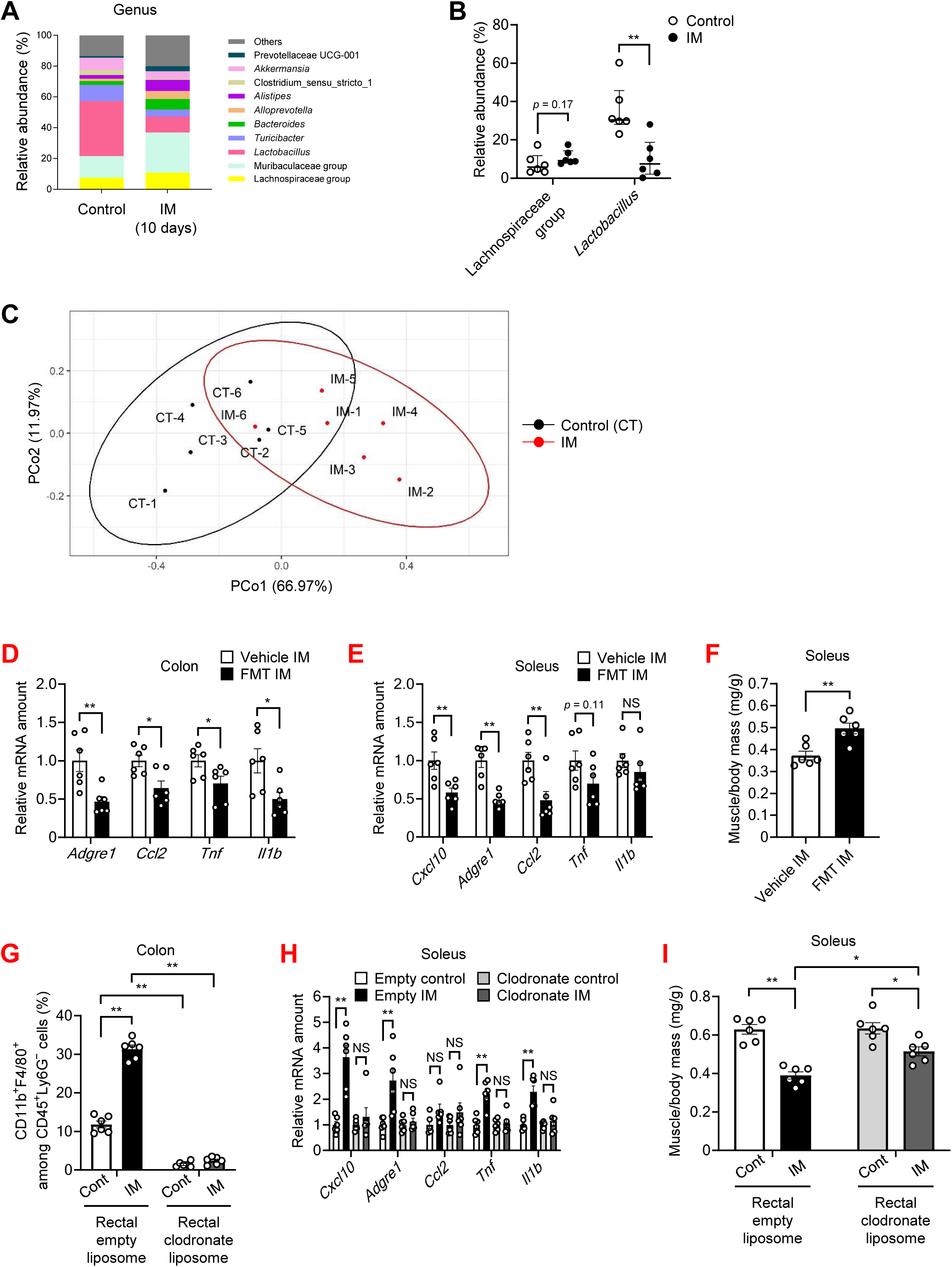
The gut microbiota and intestinal macrophages contribute to inflammation and atrophy of skeletal muscle induced by immobilization. (A–C) Gut microbial composition at the genus level determined by analysis of bacterial 16S rRNA genes (A), relative abundance of the Lachnospiraceae group and *Lactobacillus* species (B), and corresponding principal coordinate analysis (C) for control mice or mice subjected to cast-immobilization for 10 days (*n* = 6 mice). (D–F) RT-qPCR analysis of inflammation-related gene expression in the colon (D) and soleus (E), and ratio of soleus muscle mass to body mass (F) of cast-immobilized mice subjected to fecal microbiota transplantation (FMT) or vehicle by oral gavage daily for 10 days (*n* = 6 mice). (G–I) The percentage of CD11b^+^F4/80^+^ cells among CD45^+^Ly6G^−^ cells in the colon (G), RT-qPCR analysis of inflammation-related gene expression in soleus (H), and ratio of soleus muscle mass to body mass (I) of mice subjected to intrarectal injection of empty liposome or clodronate liposome every other day from 4 days before immobilization, with or without cast-immobilization for 10 days (*n* = 6 mice). Quantitative data are presented as means ± SEM (D–I) or medians with interquartile range (B). \**p* < 0.05, \*\**p* < 0.01, NS by the Mann-Whitney *U* test (B), the unpaired Student’s *t* test (D–F), or two-way ANOVA with the Bonferroni post hoc test (G–I).

Furthermore, limb immobilization by motor nerve denervation resulted in changes to the gut microbiota similar to those induced by cast-fixation (Figures S10D and S10E).

Fecal microbiota transplantation from non–cast-immobilized control mice into cast-immobilized mice attenuated colon and skeletal muscle inflammation and mitigated the loss of muscle mass induced by cast-fixation (Figures 4D–4F). Pseudosterilization of the intestine by oral administration of antibiotics also attenuated muscle inflammation and atrophy induced by cast-fixation (Figures S11A and S11B). Furthermore, intrarectal administration of clodronate liposomes inhibited the accumulation of macrophages in the colon (Figure 4G), and attenuated the upregulation of inflammation-related genes in muscle and the decrease in muscle mass induced by immobilization (Figures 4H and 4I).

These results thus implicate that alterations in the gut microbiota and intestinal macrophage activation play a critical role in mediating skeletal muscle inflammation and atrophy induced by prolonged limb immobilization.

### A gut microbial metabolite derived from linoleic acid inhibits intestinal and muscle inflammation as well as muscle atrophy induced by limb immobilization

We next examined whether the prevention of intestinal inflammation might ameliorate the loss of skeletal muscle induced by prolonged limb immobilization. We focused on gut microbial metabolites derived from linoleic acid (LA), which have been shown to exert various biological effects including the prevention of drug-induced colitis.^14–16^ Analysis of various LA-derived fatty acid metabolites (Figure S12A) in feces by mass spectrometry revealed that limb immobilization by cast-fixation for 10 days reduced the abundance of LA and 13-hydroxy-*cis*-9-octadecenoic acid (HYD) and that limb denervation for 10 days reduced that of HYD and 10-hydroxy-*cis*-12-octadecenoic acid (HYA) as well as that of metabolites derived from these two fatty acids including 13-oxo-*cis*-9-octadecenoic acid (KetoD), 10,13-dihydroxy-octadecanoic acid (HYE), 10-oxo-*cis*-12-octadecenoic acid (KetoA), and 10-oxo-*trans*-11-octadecenoic acid (KetoC) (Figures S12B and S13A), indicating that limb immobilization altered LA metabolism in the intestine.

Given that the abundance of LA, together with that of HYA and HYD, both of which are directly derived from LA, was decreased during limb immobilization, we fed mice with normal chow or chow containing these fatty acids for 10 days and then subjected them to limb immobilization. Dietary supplementation with LA or either of the LA metabolites did not influence chow intake (Figure S13B). Oral administration of HYA, but not that of LA or HYD, attenuated the reduction in crypt depth and the accumulation of F4/80-positive cells in the colon (Figures 5A–5C) as well as the shortening of the colon (Figure S14A) induced by cast-fixation. Furthermore, treatment with HYA inhibited the upregulation of inflammation-related gene expression (Figure S14B) and the downregulation of claudin-1 protein (Figure S14C) in the colon triggered by prolonged limb immobilization. Gene ontology (GO) analysis of RNA-seq data for the colon revealed that the top 20 processes enriched among genes whose expression was upregulated by immobilization include various inflammatory cell–related categories (Figure S15A) and that treatment of limb-immobilized mice with HYA attenuated the expression of genes in these categories (Figure S15B). In addition, HYA, but not LA or HYD, prevented the immobilization-induced increase in the plasma concentration of LPS (Figure S15C).

**Figure 5.**
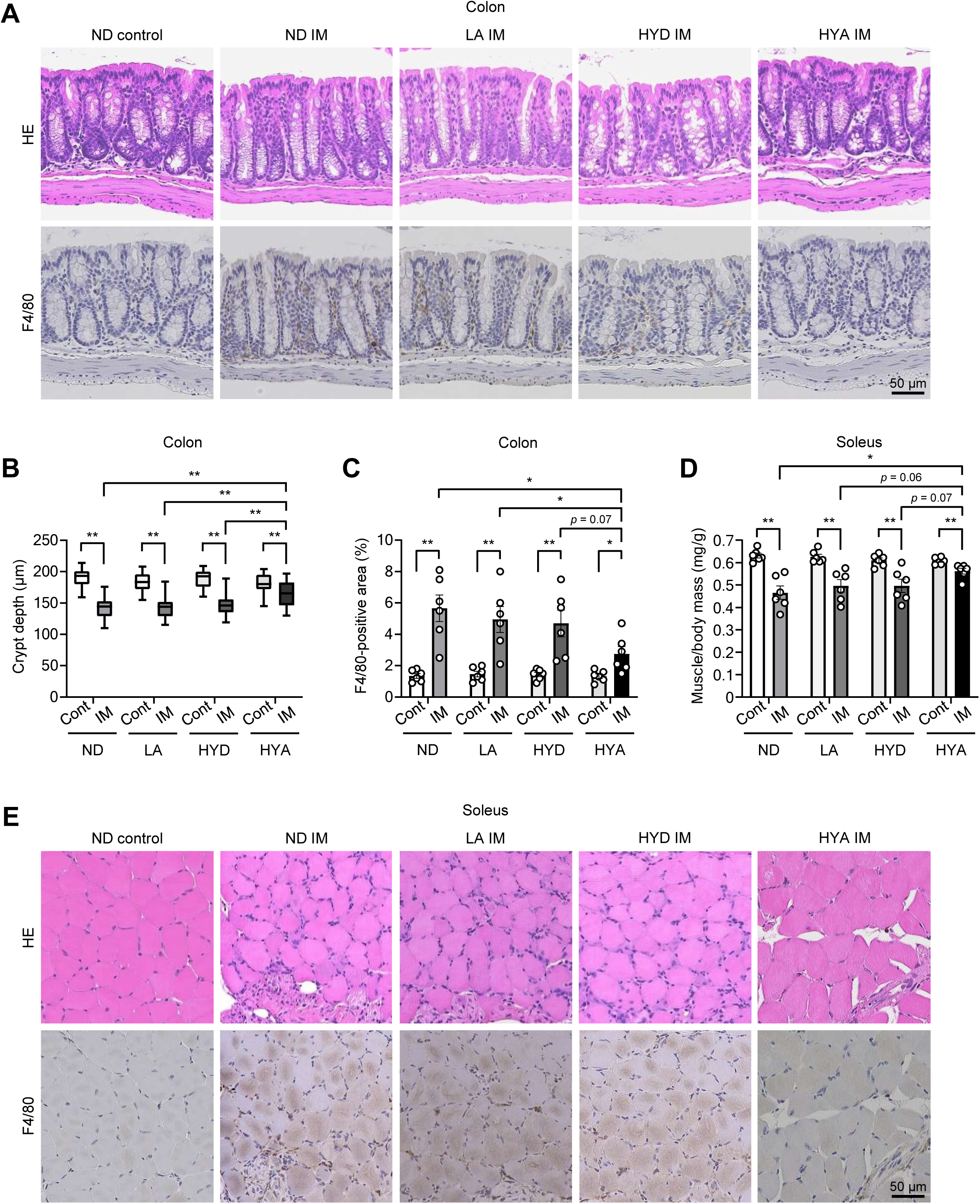
A gut microbial metabolite derived from LA inhibits intestinal and muscle inflammation as well as muscle atrophy induced by limb immobilization. (A–C) Hematoxylin-eosin and F4/80 immunohistochemical staining (A) for determination of crypt depth (B) and the F4/80-positive area (C) in the colon of mice fed a normal diet (ND) or a diet containing LA, HYD, or HYA and subjected (or not) to cast-fixation of the hind limbs for 10 days (*n* = 6 mice). The depth of 100 crypts pooled from four mice was measured and averaged for each condition in (B). Scale bar (A), 50 µm. (D and E) Ratio of muscle mass to body mass (D) as well as hematoxylin-eosin and F4/80 immunohistochemical staining (E) for the soleus of mice (*n* = 6) treated as in (A) to (C). Scale bar (E), 50 µm. Quantitative data are presented as means ± SEM (C and D) or medians with interquartile range (B). \**p* < 0.05, \*\**p* < 0.01 (two-way ANOVA with the Bonferroni post hoc test).

Oral administration of HYA, but not that of LA or HYD, also attenuated the decline in muscle mass (Figure 5D) as well as the reduction in muscle fiber area and the accumulation of F4/80-positive cells in soleus muscle (Figure 5E, Figures S16A and S16B) induced by limb immobilization. Furthermore, HYA treatment suppressed the immobilization-induced upregulation of inflammation-related gene expression in skeletal muscle (Figure S16C) while the changes in the gut microbiota caused by immobilization was unaltered by this treatment (Figure S16D). Together, these data indicated that oral administration of HYA ameliorated intestinal inflammation independently of improving dysbiosis, which in turn resulted in attenuation of inflammation and atrophy of skeletal muscle, in mice subjected to limb immobilization.

### Sympathetic nerve activation contributes to intestinal inflammation induced by limb immobilization

We next investigated the mechanism by which limb immobilization triggers intestinal inflammation. We examined the possible role of the nervous system—in particular, the sympathetic nervous system—given that sympathetic nerves regulate immune responses in various tissues including the intestine.^17,18^

Given that tissue catecholamine concentrations do not necessarily reflect sympathetic neuronal activity, we first assessed norepinephrine (NE) turnover, which is commonly used as an index of sympathetic nerve-related catecholamine dynamics.^19,20^ The NE content of tissue was thus determined before and after the intraperitoneal administration of α-methyl-*p*-tyrosine (α-MT), an inhibitor of catecholamine synthesis. As expected, α-MT treatment reduced NE content in all parts of the digestive tract examined (Figure 6A). Whereas the extent of the decrease in NE content induced by α-MT was similar for the stomach and the duodenum between control mice and mice subjected to cast-immobilization for 10 days, that for the proximal or distal small intestine or for the colon was greater in immobilized mice (Figures 6A and 6B), indicating that NE turnover was selectively augmented in the lower intestine of the immobilized animals. We also evaluated the levels of NE and its primary metabolite, DHPG (3,4-dihydroxyphenylglycol) simultaneously to assess NE dynamics, and found that the DHPG/NE ratio was markedly increased in the intestinal tissues, except for the stomach, in immobilized mice (Figure S17A), suggesting enhanced norepinephrine metabolic dynamics in the lower intestine of immobilized mice.

**Figure 6.**
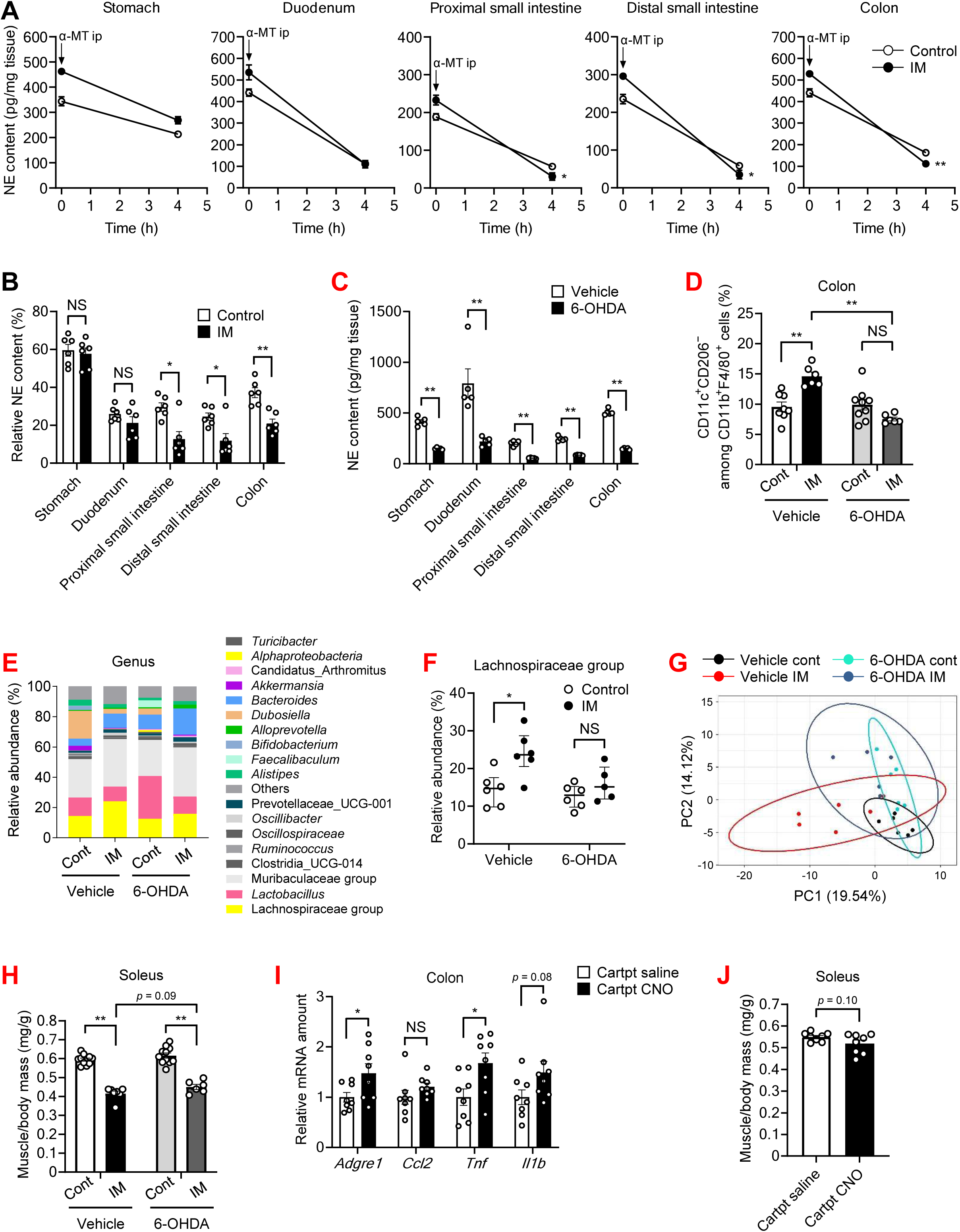
Sympathetic nerve activation contributes to intestinal inflammation and muscle atrophy induced by limb immobilization. (A and B) NE content (A) and relative NE content (at 4 h versus 0 h) (B) for the stomach, duodenum, proximal and distal small intestine, and colon of control or cast-immobilized (10 days) mice at 0 and 4 h after intraperitoneal (ip) injection of α-MT (*n* = 6 mice). (C–H) NE content for the stomach, duodenum, proximal and distal small intestine, and colon (*n* = 5 mice) (C), the percentage of CD11c^+^CD206^−^ cells among CD11b^+^F4/80^+^ cells in the colon (Vehicle control, *n* = 8; Vehicle IM, *n* = 6; 6-OHDA control, *n* = 9; and 6-OHDA IM, *n* = 6 mice) (D), gut microbial composition at the genus level (Vehicle control, *n* = 6; Vehicle IM, *n* = 6; 6-OHDA control, *n* = 6; and 6-OHDA IM, *n* = 5 mice) (E), relative abundance of the Lachnospiraceae group (Vehicle control, *n* = 6; Vehicle IM, *n* = 6; 6-OHDA control, *n* = 6; and 6-OHDA IM, *n* = 5 mice) (F), corresponding principal component analysis (Vehicle control, *n* = 6; Vehicle IM, *n* = 6; 6-OHDA control, *n* = 6; and 6-OHDA IM, *n* = 5 mice) (G), and ratio of soleus muscle mass to body mass (Vehicle control, *n* = 11; Vehicle IM, *n* = 7; 6-OHDA control, *n* = 11; and 6-OHDA IM, *n* = 6 mice) (H) of mice subjected to intraperitoneal injection of 6-OHDA or vehicle daily for 3 days, followed (or not) by cast-immobilization for 10 days starting 5 days after the final injection. (I and J) RT-qPCR analysis of inflammation-related gene expression in the colon (I) and ratio of soleus muscle mass to body mass (J) of *Cartpt-Cre* mice injected with AAV8-*hSyn-DIO-hM3D(Gq)-mCherry* into the lower thoracic spinal cord, followed 3 weeks later by intraperitoneal administration of CNO or saline daily for 7 days (*n* = 8 mice). Quantitative data are presented as means ± SEM (A–D and H–J) or medians with interquartile range (F). \**p* < 0.05, \*\**p* < 0.01, NS by the Mann-Whitney *U* test (F), the unpaired Student’s *t* test (B, C, I, and J), or two-way ANOVA with the Bonferroni post hoc test (A, D, and H).

To test the functional involvement of sympathetic nerves, we performed chemical sympathectomy using 6-hydroxydopamine (6-OHDA). 6-OHDA treatment reduced NE content of the intestinal tissues (Figure 6C) and inhibited the increase in the proportion of pro-inflammatory macrophages in the colon and the increase in the abundance of the Lachnospiraceae group, as well as the decline in skeletal muscle mass induced by immobilization (Figures 6D–6H). In addition, we selectively activated sympathetic neurons projecting to intestinal tissues by targeting *Cartpt*-expressing spinal sympathetic preganglionic neurons, which were recently shown to project selectively to the intestine.^21^ Mice with intestinal tissue-specific sympathetic nerve activation showed the upregulation of inflammation-related genes in the colon and a tendency toward reduced skeletal muscle mass (Figures 6I and 6J). These results thus indicated that activation of sympathetic nerves that selectively project to intestinal tissues contributes to intestinal inflammation, dysbiosis, and muscle atrophy induced by limb immobilization.

RNA-seq analysis of the colon revealed that, among genes encoding adrenergic receptors, the gene for the β2-adrenergic receptor (*Adrb2*) was most markedly upregulated by limb immobilization, and this upregulation was attenuated by dietary intake of HYA (Figure 7A). Expression of the gene for the α2a-adrenergic receptor (*Adra2a*) showed a similar pattern of changes, albeit to a lesser extent compared with the changes apparent for *Adrb2* expression. Treatment of mice with α-MT or the β2-adrenergic receptor antagonist propranolol, but not that with the α-adrenergic receptor antagonist phentolamine, inhibited the upregulation of inflammation-related genes in the intestine (Figure 7B) and skeletal muscle (Figure S17B). Furthermore, treatment of mice with the β2-adrenergic receptor agonist clenbuterol induced alterations in the gut microbiota that closely resembled those observed under cast immobilization (Figures 7C–7E). These results suggested that sympathetic nerve activation contributes to immobilization-induced dysbiosis and inflammation of the intestine and skeletal muscle via the activation of Adrb2 signaling. Treatment with propranolol or α-MT did not affect immobilization-induced muscle atrophy, however (Figure S17C). Given that Adrb2 in myofibers plays an important role in muscle hypertrophy,^22,23^ an antiatrophic effect of these inhibitors mediated by attenuation of Adrb2 signaling in the intestine might have been counteracted by an antihypertrophic effect mediated by attenuation of Adrb2 signaling in muscle.

**Figure 7.**
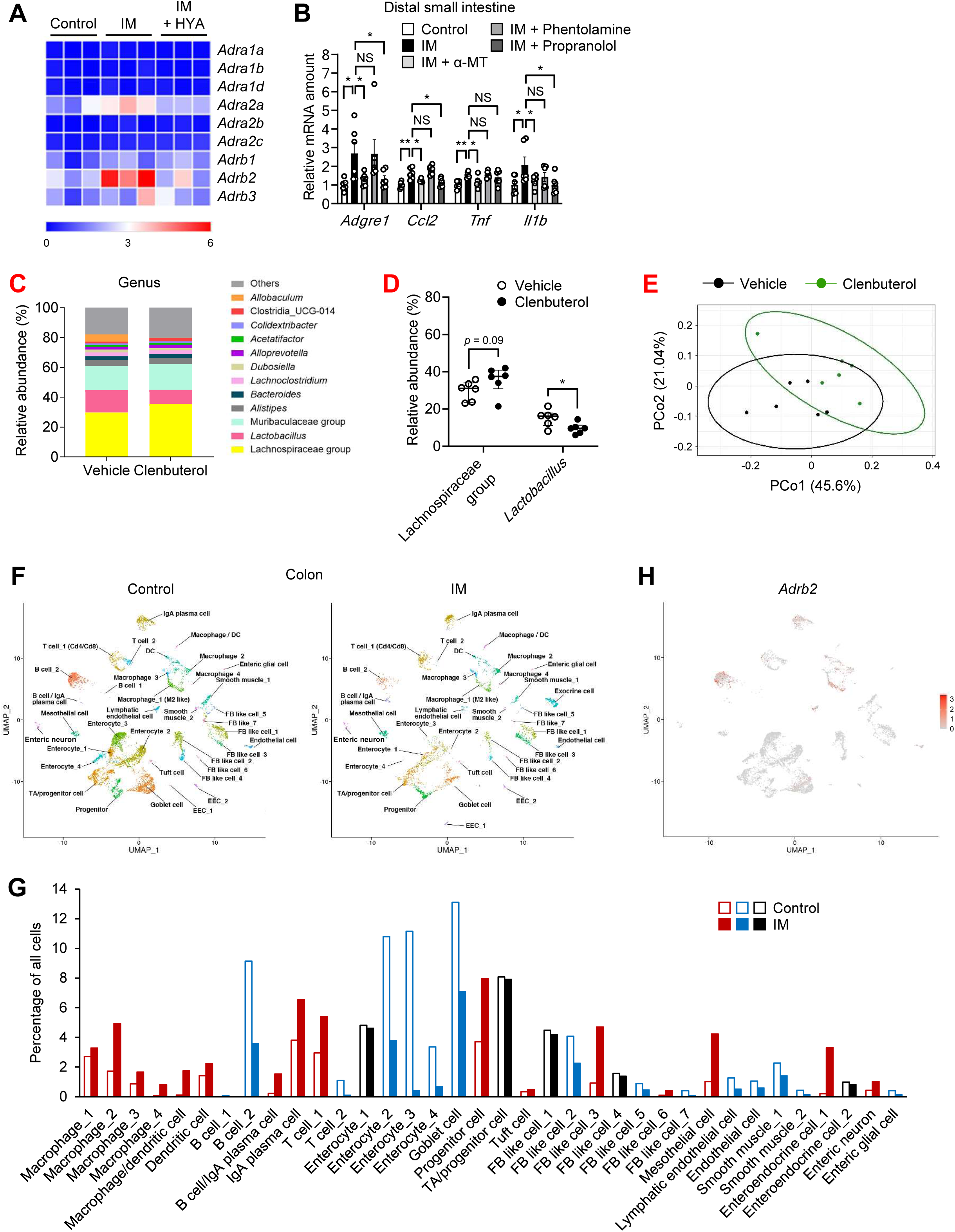
Sympathetic nerve activation contributes to immobilization-induced intestinal inflammation and dysbiosis via the activation of Adrb2 signaling. (A) Heat map for the expression of adrenergic receptor genes in the colon of mice fed a normal diet or a diet containing HYA and subjected (or not) to cast-immobilization for 10 days (*n* = 3 mice). (B) RT-qPCR analysis of inflammation-related gene expression in the distal small intestine of mice subjected to intraperitoneal injection of α-MT, phentolamine, propranolol, or vehicle daily for 10 days with or without concomitant cast-immobilization (*n* = 6 mice). (C–E) Gut microbial composition at the genus level (C), relative abundance of the Lachnospiraceae group and *Lactobacillus* species (D), and corresponding principal coordinate analysis (E) of mice subjected to intraperitoneal injection of clenbuterol or vehicle daily for 10 days (*n* = 6 mice). (F) Uniform manifold approximation and projection (UMAP) representation of 37 cell clusters identified by scRNA-seq analysis of the colon for control mice or mice subjected to cast-immobilization for 10 days. DC, dendritic cell; EEC, enteroendocrine cell; FB, fibroblast. (G) Percentage of the indicated cell clusters among all cells. Red or blue squares indicate cell clusters for which cast-immobilization increased or decreased the cell ratio by >20%, respectively. (H) Feature plot of *Adrb2* expression in the cell clusters of the colon. Quantitative data are presented as means ± SEM (B) or medians with interquartile range (D). \**p* < 0.05, \*\**p* < 0.01, NS by the Mann-Whitney *U* test (D) or two-way ANOVA with the Bonferroni post hoc test (B).

To identify cell types that mediate Adrb2 signaling in the intestine, we performed single-cell (sc) RNA-seq analysis of the colon. The single-cell fixed RNA profiling method^24^ allowed us to concomitantly analyze various cell types—including epithelial, stromal, and immune cells—while maintaining their transcriptional characteristics, leading to the identification of 37 cell clusters (Figures 7F and Figure S17D). Genes that characterize each cell cluster are listed in Table S1. Cast-immobilization of hind limbs for 10 days increased the population size for various immune cell clusters including all four macrophage clusters, a macrophage/dendritic cell cluster, a dendritic cell cluster, a B cell/IgA plasma cell cluster, an IgA plasma cell cluster, and one of the two T cell clusters (Figure 7G).

*Adrb2* mRNA was most abundant in two of the macrophage clusters (Macrophage_1 and Macrophage_2), one of the two B cell clusters (B cell_2), and a lymphatic endothelial cell cluster (Figure 7H and Figure S18). Among these cell types, only the population of macrophages was increased in response to limb immobilization (Figure 7G). We therefore focused on this cell type and generated macrophage-specific Adrb2 knockout (Mφ-Adrb2KO) mice. The immobilization-induced increase in population size was attenuated for three of the four macrophages clusters (Macrophage_2, _3, and _4), which express *Itgam* (Cd11b), *Fcer1g*, or *Cd14* and therefore likely correspond to pro-inflammatory macrophages, in Mφ-Adrb2KO mice (Figure 8A and Table S1). In contrast, *Adrb2* ablation increased the population size for the Macrophage_1 cluster, which expresses the anti-inflammatory macrophage markers *Cd163* and *Mrc1*,^25^ in the colon of mice subjected to limb immobilization (Figure 8A and Table S1). Consistent with these findings, flow cytometry revealed that the limb immobilization–induced increase in the abundance of pro-inflammatory macrophages in the colon was prevented in Mφ-Adrb2KO mice (Figures 8B and 8C, Figures S19A and S19B). Moreover, the reduction in crypt depth and the accumulation of F4/80-positive cells (Figures 8D–8F) as well as the upregulation of *Ccl2* expression (Figure S19C) in the colon induced by limb immobilization were also all attenuated in Mφ-Adrb2KO mice.

**Figure 8.**
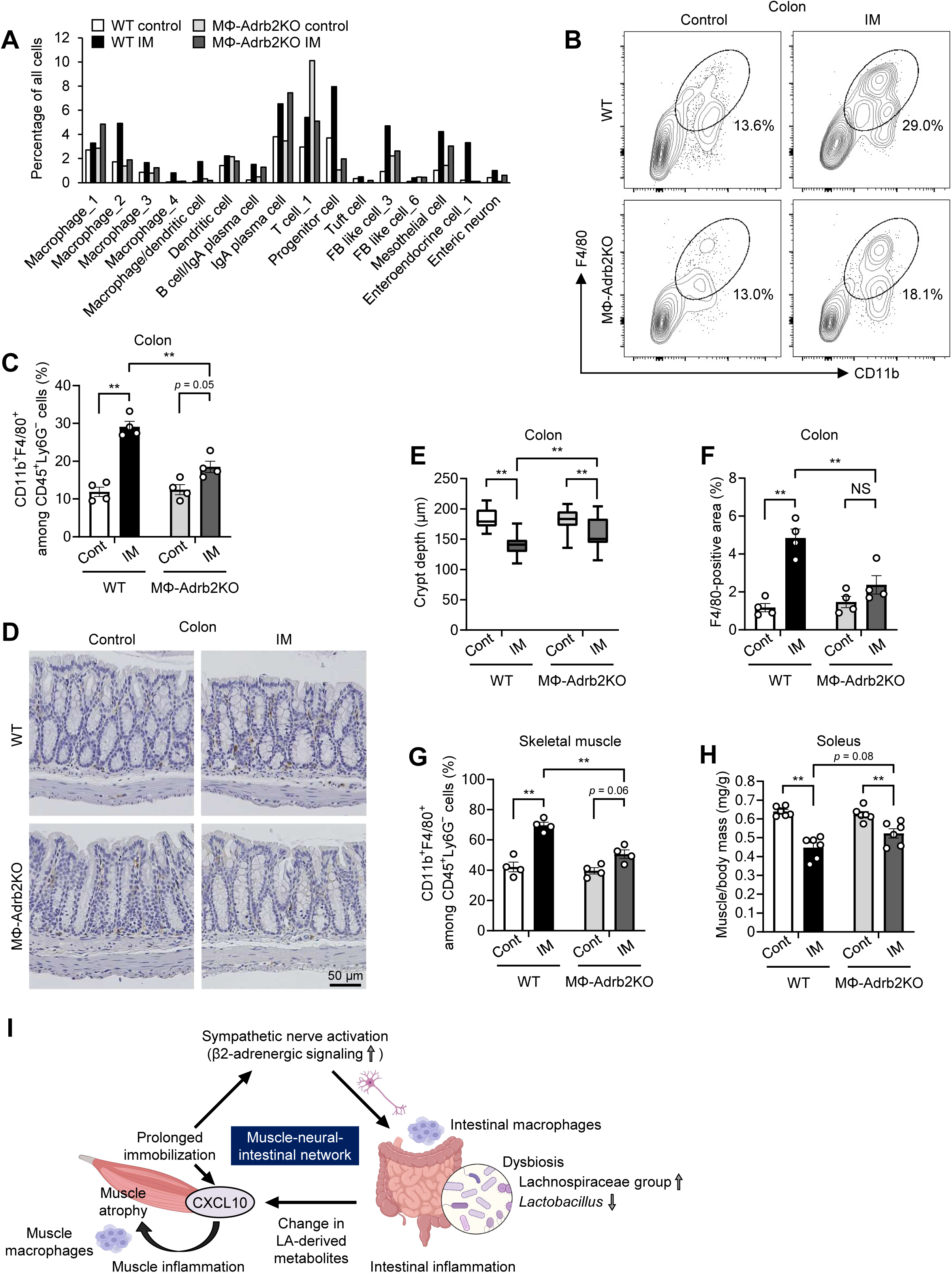
Prevention of β2-adrenergic signaling in macrophages attenuates intestinal and muscle inflammation induced by prolonged limb immobilization. (A–H) Percentage of the indicated cell clusters among all cells in the colon as determined by scRNA-seq analysis (A); representative flow cytometric analysis (B) and the percentage (*n* = 4 mice) (C) of CD11b^+^F4/80^+^ cells among CD45^+^Ly6G^−^ cells in the colon; F4/80 immunohistochemical staining of the colon (D) for determination of crypt depth (*n* = 4 mice) (E) and the F4/80-positive area (*n* = 4 mice) (F); the percentage of CD11b^+^F4/80^+^ cells among CD45^+^Ly6G^−^ cells in hind limb skeletal muscle (*n* = 4 mice) (G); and the ratio of soleus muscle mass to body mass (*n* = 6 mice) (H) for wild-type (WT) or macrophage-specific Adrb2 KO (MΦ-Adrb2KO) mice subjected (or not) to cast-immobilization for 10 days. The depth of 100 crypts pooled from four mice was measured and averaged for each condition in (E). Scale bar (D), 50 µm. (I) Proposed role of a skeletal muscle–sympathetic nerve–intestine network in immobilization-induced muscle inflammation and atrophy. Quantitative data are presented as means ± SEM (C and F–H) or medians with interquartile range (E). \*\**p* < 0.01, NS (two-way ANOVA with the Bonferroni post hoc test).

The accumulation of macrophages and the increase in the proportion of pro-inflammatory macrophages in skeletal muscle induced by prolonged limb immobilization were also inhibited, whereas the decrease in the proportion of anti-inflammatory macrophages was unaffected, in Mφ-Adrb2KO mice (Figure 8G, Figures S19D–S19F). The decline in skeletal muscle mass induced by immobilization tended to be attenuated in Mφ-Adrb2KO mice (Figure 8H). Collectively, these results suggested that Adrb2 signaling in intestinal macrophages plays a key role in skeletal muscle inflammation induced by limb immobilization.

## Discussion

We have here shown that the pathology of skeletal muscle associated with limb immobilization transitions from catabolism to inflammation, with the latter process being regulated by CXCL10 (Figure 8I). Limb immobilization results in changes to the gut microbiota and intestinal inflammation that subsequently contribute to the development of muscle inflammation. Furthermore, the onset of intestinal inflammation is triggered by sympathetic nerve activation and Adrb2 signaling in macrophages. Our results have thus uncovered a muscle-neural-intestinal network that underlies the development of muscle atrophy in response to limb immobilization.

Whereas changes to the intestinal environment contribute to various physiological and pathological processes,^26,27^ our finding that limb immobilization for a relatively short time (∼10 days) altered the gut microbiota and intestinal environment was unexpected. Our observation that the release of immobilization returned the gut microbiota to its original state indicates that the initial effect was primarily induced by immobilization via an intrinsic mechanism. Whereas extrinsic factors, including the ingestion of food or drugs, influence the gut microbiota, little is known about the intrinsic mechanisms by which the host modulates the gut microbial population. In addition to its effects on immune cell clusters, limb immobilization increased the population size for a progenitor cell cluster, a tuft cell cluster, two of the seven fibroblast-like cell clusters (FB like cells_3 and _6), a mesothelial cell cluster, one of the two enteroendocrine cell clusters (Enteroendocrine cell_1), and an enteric neuron cluster in the intestine. Changes in population size and function of these cell clusters may also have contributed to the immobilization-induced alterations in the intestinal environment and gut microbiota.

Metabolites produced by the gut microbiota play an important role in symbiosis with the host. HYA is a recently discovered LA-derived metabolite produced by gut bacteria and has been detected in human plasma.^14^ Whereas the roles of HYA and its related lipid metabolites in the maintenance of human physiological functions are not yet fully understood, evidence suggests that exogenous HYA administration may ameliorate various pathological conditions including chemical colitis, atopic dermatitis, fatty liver disease, and obesity-induced insulin resistance.^28–31^ The mechanisms by which HYA exerts such beneficial effects on these conditions, as well as on immobilization-induced muscle inflammation, remain unclear. However, HYA not only attenuates impairment of the intestinal barrier but also interacts with several host proteins including fatty acid receptors.^28^ Further elucidation of the detailed mechanisms of HYA action may contribute to the development of its clinical use.

Sympathetic nerves together with immune cells that express adrenergic receptors in the intestine have been implicated in the regulation of tissue damage and repair.^17,18^ Adrb1 and Adrb2 expressed in innate lymphoid cells of the small intestine contribute to protection of the tissue against inflammation^32^ and to tissue regeneration.^33^ The T cell_2 cluster in our analysis appears to correspond to innate lymphoid cells, with the population size for this cell cluster being decreased by limb immobilization.

Macrophages that reside in the muscular layer of the intestine exert tissue-protective actions dependent on Adrb2 signaling during bacterial infection.^34,35^ We have now shown that *Adrb2* was highly expressed in the Macrophage_1 and _2 clusters, which manifested transcriptional characteristics of anti-inflammatory and pro-inflammatory macrophages, respectively. Macrophage_1 cells therefore likely correspond to macrophages that reside in the muscular layer of the intestine.

Our findings have potential implications for drug development. Given that the safety of HYA administration in humans has been established, we have initiated a clinical trial of such treatment for individuals prone to the development of muscle atrophy as a result of limb immobilization after orthopedic surgery (clinical trial number: jRCT2051230197). Whereas antibodies to CXCL10 have been shown to be effective for the treatment of inflammatory and cardiovascular conditions in preclinical studies,^36–38^ a preventive effect of these antibodies on muscle atrophy has not been reported. Whereas several clinical trials of agents that target CXCL10 have been conducted,^39,40^ such agents have yet to be introduced into clinical practice. Given that the upregulation of *CXCL10* expression in muscle was associated with acute limb immobilization not only in the human cohort of the present study but also in a publicly available data set, clinical trials to explore the potential preventive effect of antibodies to CXCL10 on immobilization-induced muscle atrophy are warranted.

In conclusion, our study has revealed a previously unrecognized organ-organ interaction that plays a key role in the development of immobilization-induced muscle atrophy. Two different interventions, both of which are immediately applicable to clinical trials, were shown to be effective for prevention of this condition: the targeting of muscle inflammation with antibodies to CXCL10, and that of intestinal inflammation with HYA. The mechanisms by which limb immobilization signals to the sympathetic nervous system and by which Adrb2 signaling in macrophages induces intestinal inflammation remain to be elucidated. The characterization of these mechanisms may contribute to the further development of new and improved treatments for muscle atrophy.

## Acknowledgments

We thank Akira Suzuki for providing *LysM-Cre* mice, Hidenori Shimizu for gut microbiota analysis, and Chikako Aoki and Yoshika Ogawa for technical assistance. We also thank Mashito Sakai for the insightful discussions. This study was supported in part by grants from Japan Society for the Promotion of Science (KAKENHI grant 22K16397 to Y. Hirata) and Japan Agency for Medical Research and Development (AMED) Practical Research Project for Life-Style related Diseases including Cardiovascular Diseases and Diabetes Mellitus (Grant Number JP23ek0210192 to Y. Hirata, K.N., and W.O.), as well as by grants from Uehara Memorial Foundation, Nakatomi Foundation, Japan Society for the Study of Obesity, Center for Medical Transformation, Manpei Suzuki Diabetes Foundation, and Cell Science Research Foundation (all to Y. Hirata), and from Takeda Science Foundation (to W.O.).

## Author Contributions

Y. Hirata, K.N., and W.O. conceived the study and analyzed the data. K.H., Y. Hosokawa, K.U., T.I., T. Nishigaki, A.L., S.Y., Y.M., and K.M. contributed to animal and cell experiments. K.S. contributed to analysis of the public data set. T. Niikura, T.F., K. Oe, R.K., and Y. Hirota contributed to analysis of human skeletal muscle. S.H. contributed to gut microbiota and metabolome analysis. S.K. and J.H. contributed to RNA-seq analysis. G.K. contributed to the generation of genetically engineered mice. K. Ohbayashi, T.Y., and Y.I. contributed to measurement of NE turnover. C.K.G. provided additional experimental guidance. Y. Hirata and W.O. wrote the manuscript.

## Supplemental information

Document S1. Figures S1–S20 and Tables S1–S3

## Methods

### Animals

*Adrb2*-floxed mice were described previously,^41^ and *LysM-Cre* mice^42^ were obtained from The Jackson Laboratory (JAX stock #004781). These two mouse lines were crossed to obtain macrophage-specific Adrb2 KO (MΦ-Adrb2KO) mice. All experiments were performed with male mice at 10 weeks of age unless indicated otherwise. Mice were anesthetized by intraperitoneal injection of a mixture of medetomidine (0.3 mg/kg), midazolam (4.0 mg/kg), and butorphanol (5.0 mg/kg) and were then subjected either to bilateral cast-immobilization of hind limbs with the use of plastic tubes, adhesive tape, and animal bandage, or to bilateral hind limb denervation by transection of the sciatic nerve. The immobilized mice were able to move freely around their cages and to obtain food and water using their forelimbs. Animals were analyzed 3 or 10 days after cast-immobilization or denervation unless indicated otherwise. Mice were injected intraperitoneally with neutralizing antibodies to CXCL10 (MAB466, clone 134013; R&D Systems) or control IgG at 0.1 mg/body 3 days before cast-immobilization, or with α-MT (Sigma-Aldrich) at 100 mg/kg, phentolamine (Tokyo Chemical Industry) at 20 mg/kg, or propranolol (Sigma-Aldrich) at 10 mg/kg daily for 10 days concomitant with cast-immobilization, or clenbuterol (Sigma-Aldrich) at 1 mg/kg daily for 10 days. For fecal microbiota transplantation,^43^ mice were subjected to oral gavage with 200 μl supernatant of freshly collected fecal samples (50 mg/ml) or vehicle daily for 10 days. For antibiotic treatment, mice were subjected to oral gavage with a cocktail of ampicillin (5 mg/ml), gentamicin (5 mg/ml), neomycin (5 mg/ml), metronidazole (5 mg/ml), and vancomycin (2.5 mg/ml) daily from 4 days before immobilization, with or without cast-immobilization for 10 days. To deplete macrophages in the colon,^44^ mice were subjected to intrarectal injection of empty liposome or clodronate liposome (Encapsula NanoSciences) at 500 μg/100 μl with flexible catheter every other day from 4 days before immobilization, with or without cast-immobilization for 10 days. Diets containing LA, HYD, or HYA (each at 1%) were obtained from Noster and were administered to mice beginning 10 days before immobilization. For chemical sympathectomy,^45^ mice were subjected to intraperitoneal injection of 6-OHDA (Sigma-Aldrich) at 80 mg/kg or vehicle (0.2% ascorbic acid, Tocris Bioscience) daily for 3 days, with or without cast-immobilization for 10 days starting 5 days after the final injection. *Cartpt-Cre* mice were described previously.^21^ AAV8-*hSyn-DIO-hM3D(Gq)-mCherry* was injected into the lower thoracic spinal cord, and three weeks later, 2 mg/kg clozapine-N-oxide (CNO) or saline was administered intraperitoneally daily for 7 days. The separation of blood cells from non–blood cells of skeletal muscle was performed with the use of biotin-conjugated antibodies to mouse CD45.2 (109804, BioLegend), microbeads conjugated with antibodies to biotin (130-090-485, Miltenyi Biotec), and an autoMACS Pro Separator (Miltenyi Biotec). The concentration of LPS or CXCL10 in the plasma or muscle tissue was measured using the LAL endotoxin assay kit (GenScript) or the mouse CXCL10 DuoSet ELISA kit (R&D Systems).

### Cell culture, RT-qPCR analysis, and immunoblot analysis

C2C12 myoblasts were maintained and induced to differentiate into myotubes as described previously,^46^ and they were treated with LPS (L3129, Sigma-Aldrich) at 1 µg/ml for 6 h. Isolation of total RNA from cultured myotubes or from mouse or human tissues as well as RT-qPCR analysis were performed as previously described,^46^ and the abundance of target mRNAs was normalized by that of 36B4 mRNA. The sequences of mouse and human PCR primers are provided in Tables S2 and S3, respectively.

Immunoblot analysis was performed with antibodies to claudin-1 (2H10D10, Invitrogen), GAPDH (M171-7, MBL), ERK (9102, Cell Signaling Technology), and Thr^202^/Tyr^204^-phosphorylated ERK (9101, Cell Signaling Technology). Uncropped immunoblots are shown in Figure S20.

### Histology and immunohistochemical staining

Skeletal muscle or colon tissue sections were stained with hematoxylin-eosin or subjected to immunohistochemical staining with antibodies to F4/80, myosin heavy chain 7 (slow-twitch), or myosin heavy chain 1 (fast-twitch). Images were acquired with a BZ-X710 fluorescence microscope (Keyence). The area of muscle fibers, depth of crypts, and F4/80-positive area were quantified for each image with the use of ImageJ software (NIH).

### Flow cytometry

Skeletal muscle or the colon was minced and digested with 0.2% collagenase type II (Worthington), the digested tissue was passed through 100-μm and 40-μm cell strainers (Falcon), and erythrocytes were eliminated from the filtrate. The remaining cells were exposed to antibodies to mouse CD16/32 (clone 93, 101302; BioLegend) before staining with antibodies to the following proteins: CD45.2 (clone 104, 109822; BioLegend), Ly6G (clone 1A8, 127606; BioLegend), F4/80 (clone BM8, 123116; BioLegend), CD11b (clone M1/70, 101224; BioLegend), CD11c (clone N418, 117308; BioLegend), and CD206 (clone C068C2, 141720; BioLegend). Dead cells were excluded by staining with propidium iodide (421301, BioLegend). Events were acquired with an LSR Fortessa X-20 instrument (BD Biosciences) and analyzed with FlowJo software (BD Biosciences).

### DNA microarray analysis

Total RNA extracted from gastrocnemius muscle of mice at 3 days after cast immobilization or of corresponding control animals was subjected to hybridization with an Affymetrix Mouse Gene 2.0 ST Array.

### RNA-seq and scRNA-seq analysis

Total RNA extracted from gastrocnemius muscle, flexor carpi ulnaris, or colon tissue was subjected to RNA-seq analysis by Macrogen or Novogene. A paired-end library was prepared from each RNA sample with the use of a TruSeq Stranded mRNA Library Preparation Kit and was sequenced with a NovaSeq 6000 system (Illumina). Data were analyzed with RaNA-Seq^47^ and DAVID Bioinformatics Resources 6.8. A Chromium Next GEM Single Cell Fixed RNA Sample Preparation Kit was used to isolate cells from colon tissue, and an scRNA-seq library was generated with a 10x Chromium system. Data analysis was performed by Genble.

### Gut microbiota and metabolome analysis

DNA was extracted from feces with the use of a QIAamp PowerFecal Pro DNA Kit (Qiagen). For analysis of 16S rRNA genes of the gut microbiota, the V3-V4 region of the gene was amplified from the extracted DNA with 341F (5′-TCGTCGGCAGCGTCAGATGTGTATAAGCGACAGCCTACGGGNGGCWGCAG-3′) and 805R (5′-GTCTCGTGGGCTCGGAGATGTGTATAAGAGACAGGACTACHVGGGTATCTA ATCC-3′) primers. Both ends of the amplified fragments were tagged with the use of a Nextera XT Index Kit v2 (Illumina), and the amplicons were then purified and sequenced with the use of an Illumina MiSeq system and MiSeq Reagent Kit v3.

Postsequencing analysis was performed with the Quantitative Insights Into Microbial Ecology 2 (QIIME2) pipeline. SILVA (version SSU138.1) was adopted as a reference sequence database for estimation of bacterial species prevalence, and gut microbiota composition was determined with OTUs, which grouped reads with a sequence identity of ≥97%. Comprehensive analysis of gut microbial lipid metabolites was performed as previously described.^48^

### Measurement of NE turnover

NE turnover was measured on the basis of the decline in tissue NE content after inhibition of catecholamine biosynthesis with α-MT as described previously.^19,20^ Mice were injected intraperitoneally with α-MT at 300 mg/kg. At 0 or 4 h after α-MT injection, the animals were decapitated and tissues were rapidly removed and weighed. Tissue samples were homogenized in 0.2 M perchloric acid containing 0.1 mM EDTA, the homogenate was centrifuged at 4°C, and NE or DHPG was purified from the resulting supernatant with activated alumina and measured by high-performance liquid chromatography with electrochemical detection. Data are expressed as picograms of NE or DHPG per milligram of tissue weight.

### Human skeletal muscle samples

Specimens of human skeletal muscle were obtained as previously described.^6^ In brief, samples were obtained from 15 individuals at an average of 8.3 ± 3.1 days after fixation surgery for a bone fracture of the arm or leg as well as from 18 control participants who underwent implant-removal surgery 6 to 12 months after fixation surgery.

### Publicly available human gene expression data analysis

For the analysis of publicly available human gene expression data, the process of data set selection was undertaken as follows. NCBI GEO and Array Express databases were searched with the terms “muscle”, “cast”, “immobilization”, and “human”. Screening of the three identified data sets resulted in the exclusion of one (GSE45462) because it included only individuals with fractures. The remaining two data sets (GSE14901 and GSE8872) underwent eligibility assessment, with GSE8872 being subsequently excluded on the basis of a small number of healthy subjects (*n* = 2). The remaining data set (GSE14901) met all the inclusion criteria and was adopted for analysis.

## Statistical analysis

Quantitative data were presented as means ± SEM or as medians with the interquartile range, and they were analyzed by the two-tailed unpaired Student’s *t* test, two-way analysis of variance (ANOVA) with Bonferroni’s post hoc test, the Mann-Whitney *U* test, or one-way ANOVA followed by Dunnett’s test or Bonferroni’s post hoc test. A *p* value of <0.05 was considered statistically significant. All statistical analysis was performed with the use of GraphPad Prism 9 (GraphPad Software).

## Study approval

All animal experiments were approved by the animal experimentation committee of Kobe University Graduate School of Medicine (approval no. P221201). Human experiments were approved by the medical ethics committee of Kobe University Graduate School of Medicine (approval no. 180059), and all subjects provided written informed consent.

## Data availability

The microarray data have been deposited in NCBI Gene Expression Omnibus (GEO) under the accession number GSE172501. The RNA-seq data have been deposited in NCBI GEO under the accession numbers GSE249515 (mouse gastrocnemius), GSE288180 (mouse flexor carpi ulnaris), and GSE249324 (mouse colon). The scRNA-seq data have been deposited in NCBI GEO under the accession number GSE265821.

---

**Figure S1.**
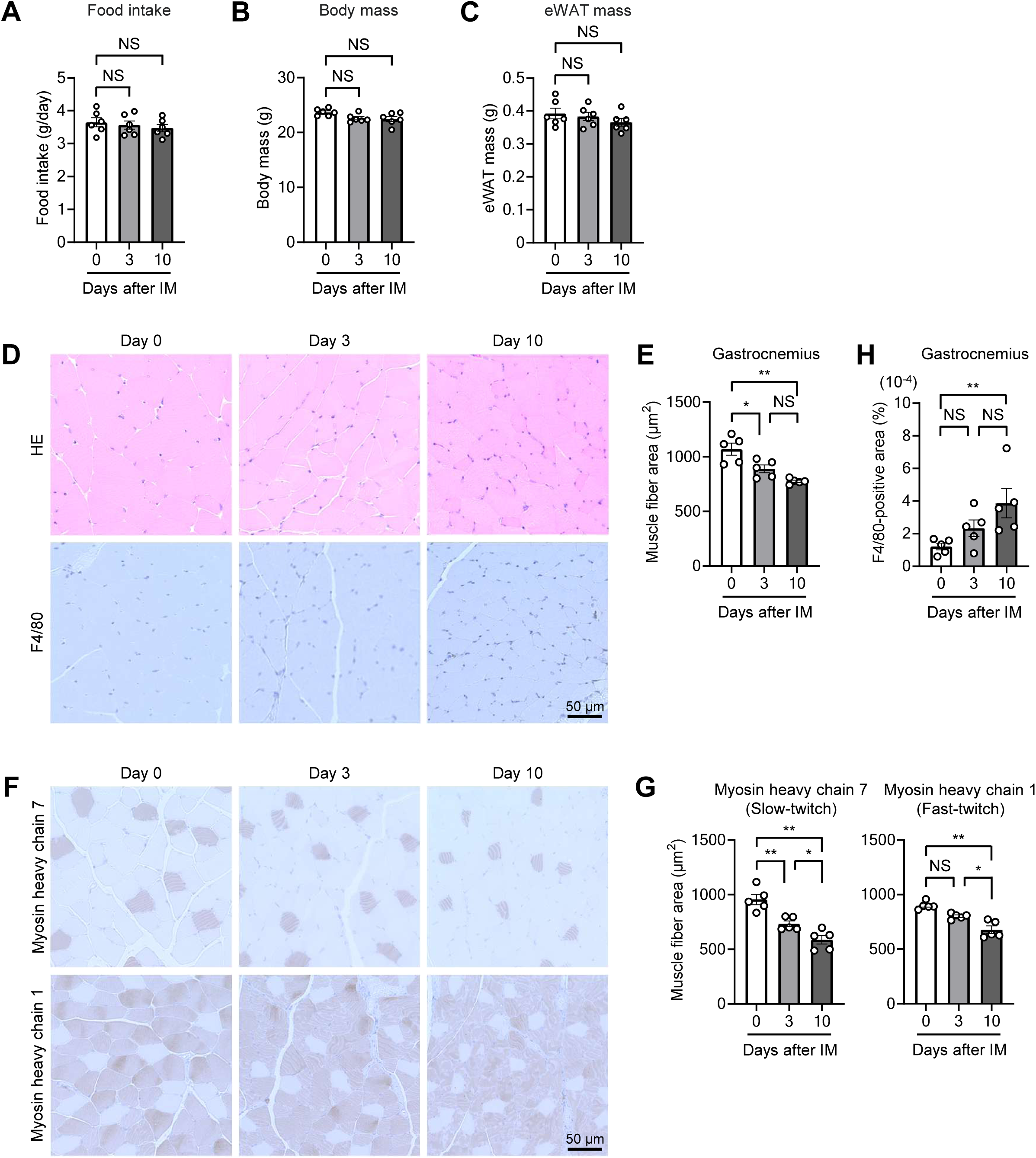
Hematoxylin-eosin staining and immunohistochemical staining in skeletal muscle of immobilized mice. (A–C) Daily food intake (A), body mass (B), and epididymal white adipose tissue (eWAT) mass (C) for mice subjected to bilateral hind limb immobilization by cast-fixation for the indicated times (*n* = 6 mice). (D and E) Hematoxylin-eosin staining and immunohistochemical staining of F4/80 (D) for determination of muscle fiber area (E) in gastrocnemius muscle of mice subjected to hind limb immobilization for the indicated times (*n* = 5 mice). The area of fibers pooled from five mice was measured and averaged for each condition in (E). Scale bar (D), 50 μm. (F and G) Immunohistochemical staining of myosin heavy chain 7 and myosin heavy chain 1 (F) for determination of muscle fiber area (G) in gastrocnemius muscle of mice subjected to hind limb immobilization for the indicated times (*n* = 5 mice). The area of fibers pooled from five mice was measured and averaged for each condition in (G). Scale bar (F), 50 μm. (H) The F4/80-positive area in gastrocnemius muscle of mice subjected to hind limb immobilization for the indicated times (*n* = 5 mice). Quantitative data are presented as means ± SEM (A–C, E, G, and H). \**p* < 0.05, \*\**p* < 0.01, NS by two-way ANOVA with the Bonferroni post hoc test.

**Figure S2.**
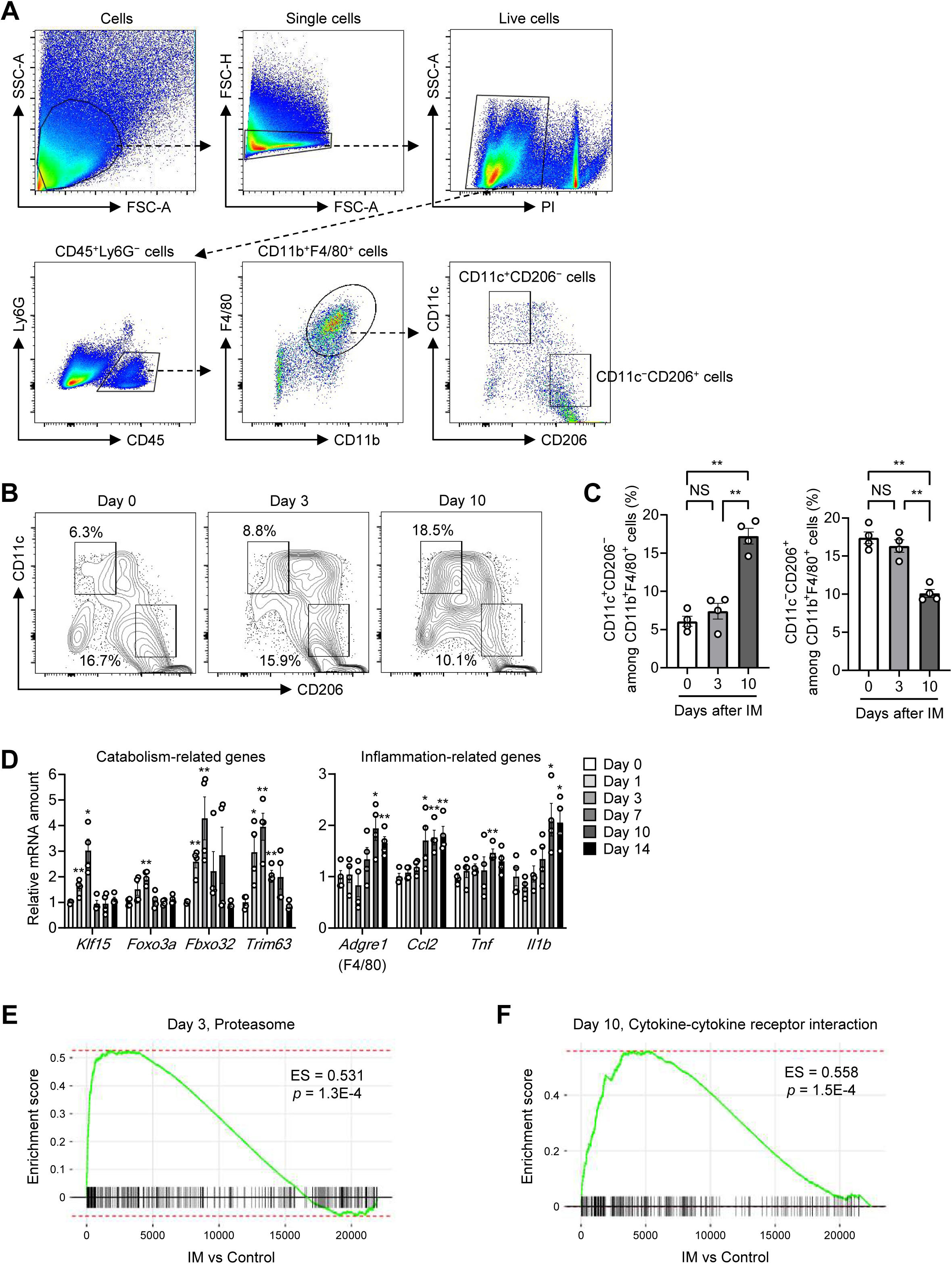
Gating strategy and immobilization-induced changes in macrophage subsets of skeletal muscle as well as GSEA for immobilized muscle. (A) Gating strategy for fluorescence-activated sorting of macrophages from skeletal muscle of immobilized mice. SSC, side scatter; FSC, forward scatter; PI, propidium iodide. (B and C) Representative flow cytometric analysis (B) and the percentage (C) of CD11c^+^CD206^−^ or CD11c^−^CD206^+^ cells in skeletal muscle of mice subjected to hind limb immobilization by cast-fixation for the indicated times (*n* = 4 mice). (D) RT-qPCR analysis of protein catabolism– or inflammation-related gene expression in soleus muscle of immobilized mice (*n* = 4 mice). (E and F) GSEA for the indicated pathways performed with RNA-seq data from gastrocnemius muscle of mice immobilized for 3 days (E) or 10 days (F). ES, enrichment score. Quantitative data are presented as means ± SEM (C and D). \**p* < 0.05, \*\**p* < 0.01, NS compared with day 0 (D) or as indicated (C) by two-way ANOVA with the Bonferroni post hoc test.

**Figure S3.**
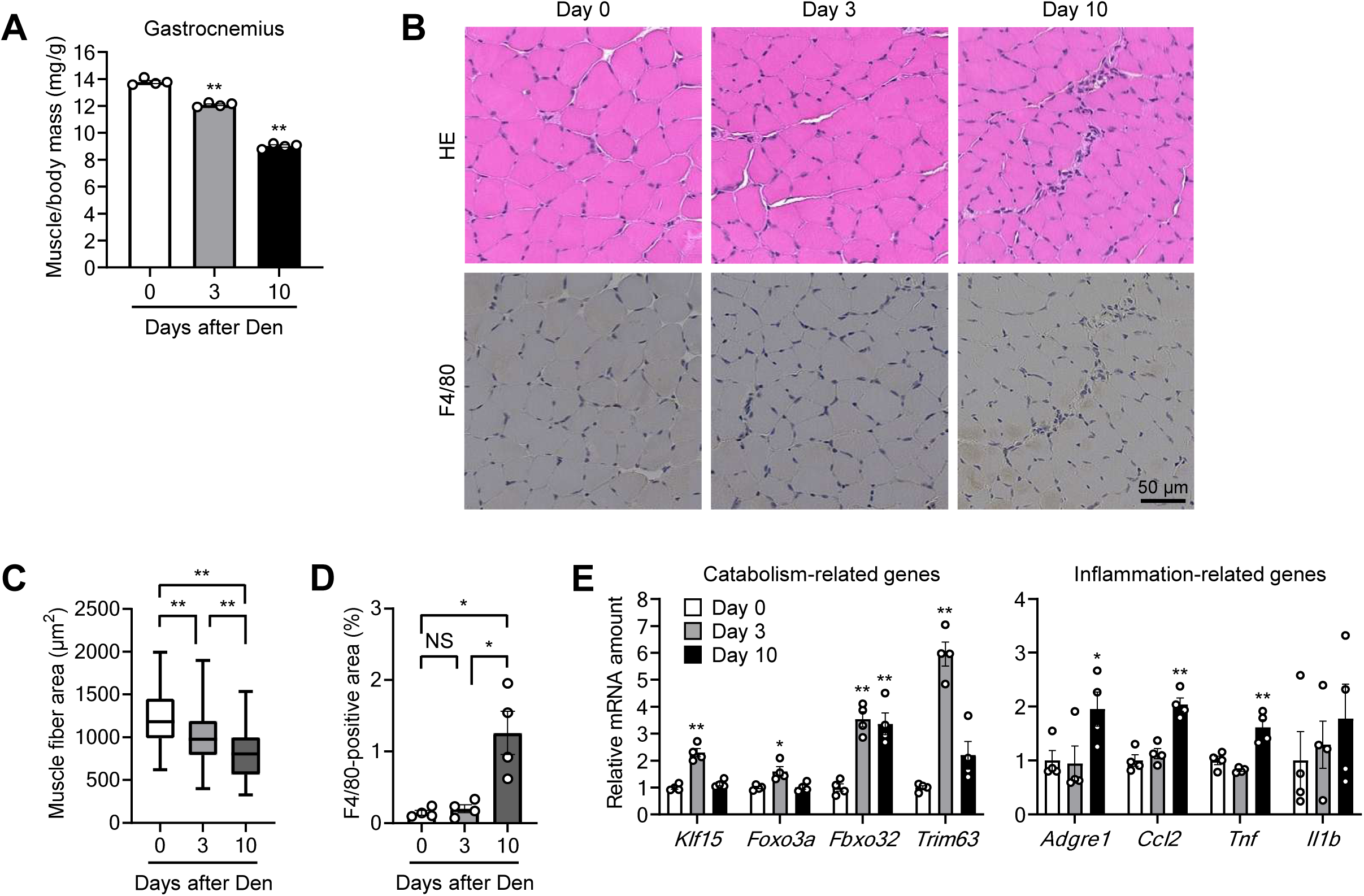
Time course of denervation-induced muscle atrophy, macrophage accumulation, and changes in gene expression. (A) Ratio of gastrocnemius muscle mass in both hind limbs to body mass for mice subjected to bilateral hind limb denervation (Den) for the indicated times (*n* = 4 mice). (B–D) Hematoxylin-eosin and F4/80 immunohistochemical staining (B) for determination of muscle fiber area (C) and the F4/80-positive area (D) in soleus muscle of denervated mice (*n* = 4 mice). The area of 800 fibers pooled from four mice was measured and averaged for each condition in (C). Scale bar (B), 50 μm. (E) RT-qPCR analysis of protein catabolism– or inflammation-related gene expression in the gastrocnemius of denervated mice (*n* = 4 mice). Quantitative data are presented as means ± SEM (A, D, and E) or medians with interquartile range (C). \**p* < 0.05, \*\**p* < 0.01, NS compared with day 0 (A and E) or as indicated (C and D) by two-way ANOVA with the Bonferroni post hoc test.

**Figure S4.**
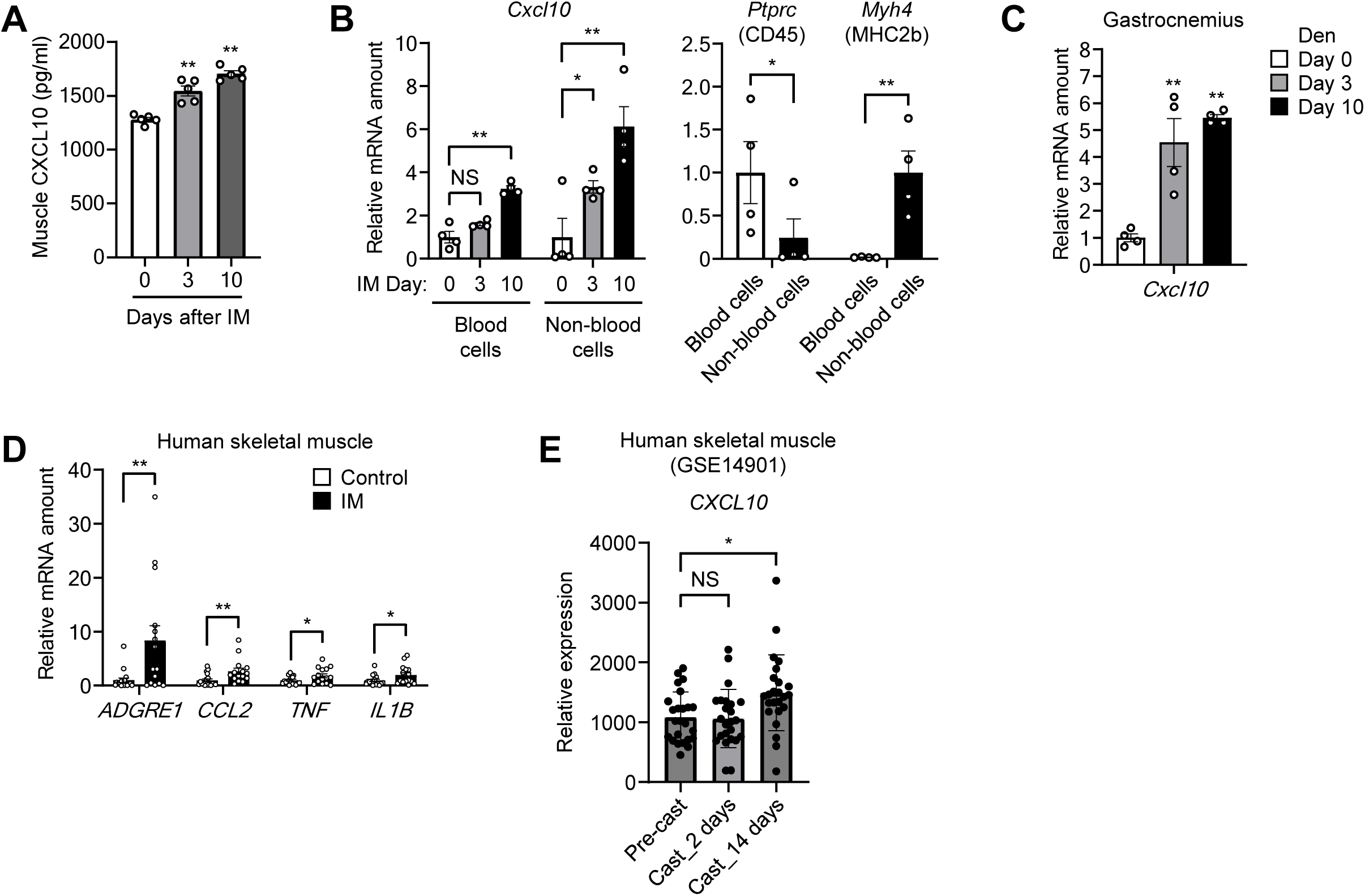
Identification of the chemokine CXCL10 as a mediator of muscle inflammation induced by limb immobilization. (A) Muscle tissue concentration of CXCL10 in cast-immobilized mice (*n* = 5 mice). (B) RT-qPCR analysis of *Cxcl10*, *Ptprc*, and *Myh4* mRNA abundance in the blood cell and non–blood cell fractions of skeletal muscle from cast-immobilized mice (*n* = 4 mice). (C) RT-qPCR analysis of *Cxcl10* mRNA in gastrocnemius of denervated mice (*n* = 4 mice). (D) RT-qPCR analysis of inflammation-related gene expression in skeletal muscle of control (*n* = 18) or immobilized (*n* = 15) human participants. (E) Relative expression of *CXCL10* in skeletal muscle of healthy human individuals subjected to cast-immobilization as determined from a publicly available data set. The relative expression level is presented as transcripts per million (TPM) for the subjects before (Pre-cast) and at two time points after (Cast_2 days and Cast_14 days) cast application. Each point represents one individual. Quantitative data are presented as means ± SEM (A–E). \**p* < 0.05, \*\**p* < 0.01, NS compared with day 0 (A and C) or as indicated (B, D, and E) by the unpaired Student’s *t* test (B and D), two-way ANOVA with the Bonferroni post hoc test (A–C), or one-way ANOVA followed by Dunnett’s test (E).

**Figure S5.**
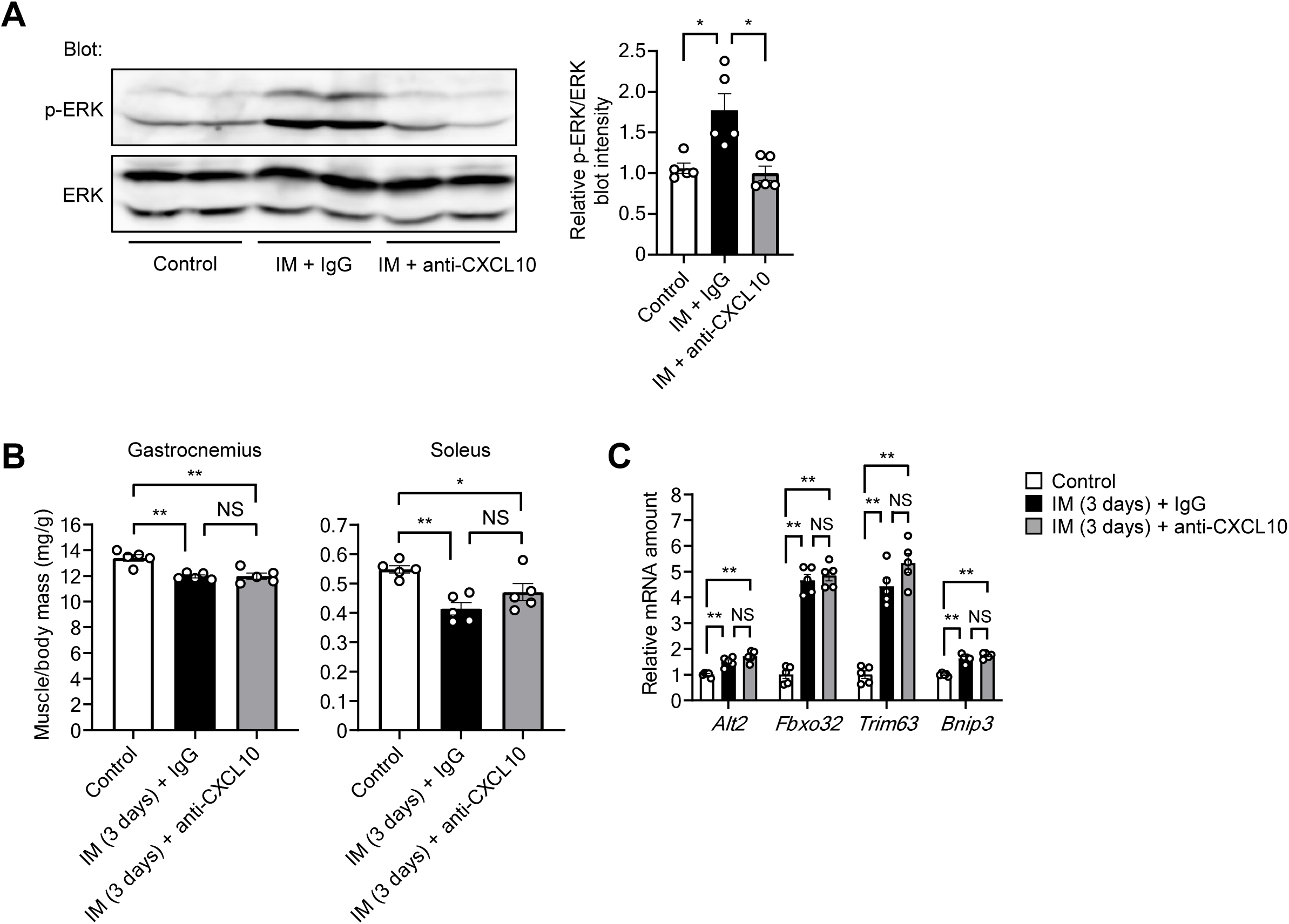
Effects of neutralizing antibodies to CXCL10 on muscle inflammation and atrophy induced by immobilization. (A) Immunoblot analysis of total and phosphorylated (p-) forms of ERK in gastrocnemius for control mice or mice subjected to intraperitoneal injection of neutralizing antibodies to CXCL10 (0.1 mg/body) or control IgG 3 days before cast-immobilization for 10 days (n = 5 mice). (B and C) Ratio of muscle mass to body mass (B) and RT-qPCR analysis of catabolism-related gene expression in gastrocnemius (C) for control mice or mice subjected to intraperitoneal injection of neutralizing antibodies to CXCL10 (0.1 mg/body) or control IgG 3 days before cast-immobilization for 3 days (*n* = 5 mice). Quantitative data are presented as means ± SEM (A–C). \**p* < 0.05, \*\**p* < 0.01, NS by two-way ANOVA with the Bonferroni post hoc test.

**Figure S6.**
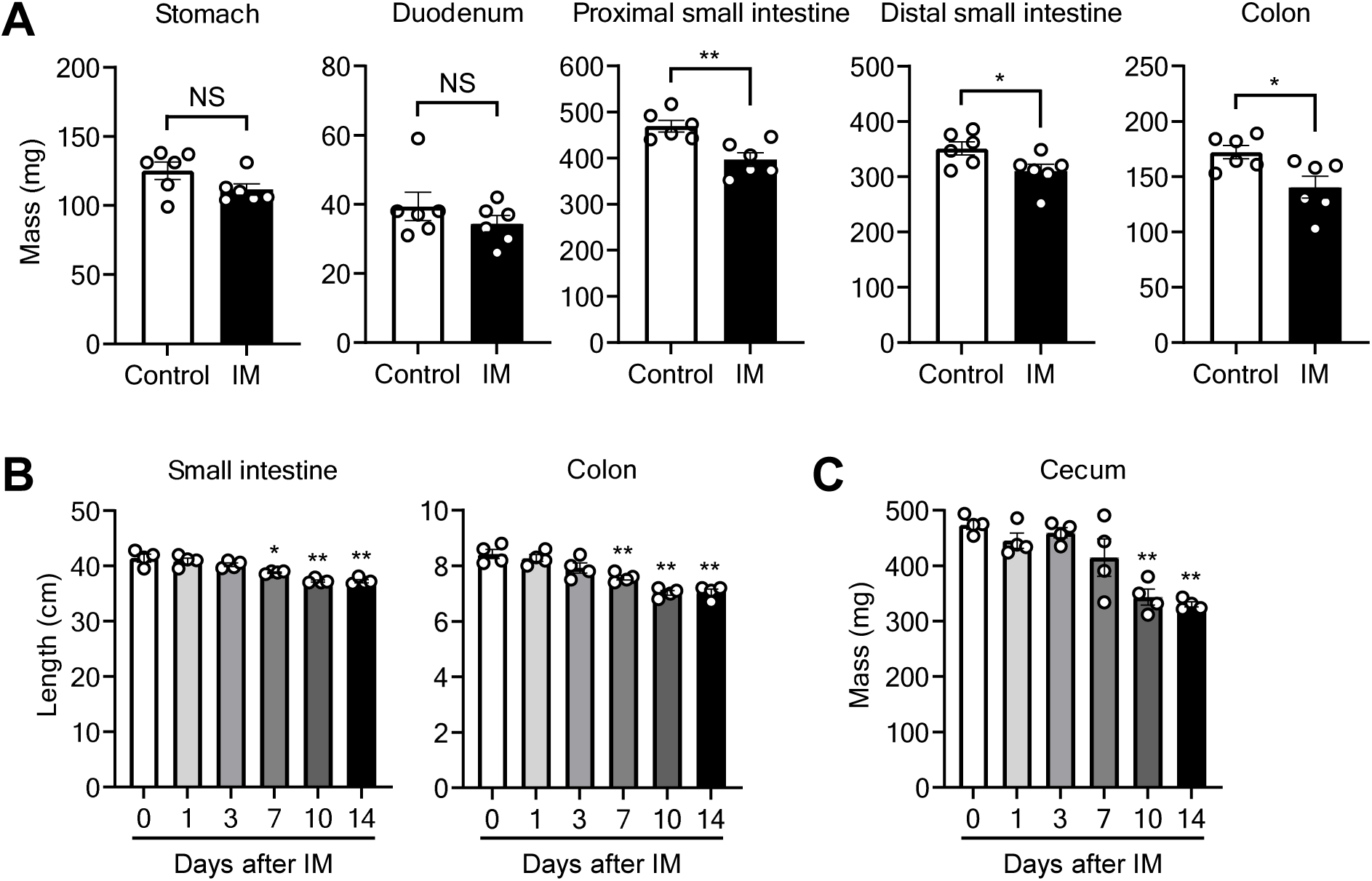
Effects of prolonged limb immobilization on intestinal tissue. (A) Mass of the stomach, duodenum, proximal and distal small intestine, and colon for control mice or mice subjected to cast-immobilization for 10 days (*n* = 6 mice). (B and C) Length of the small intestine and colon (B) as well as mass of the cecum (C) for mice subjected to cast-immobilization for the indicated times (*n* = 4 mice). Quantitative data are presented as means ± SEM (A–C). \**p* < 0.05, \*\**p* < 0.01, NS compared with day 0 (B and C) or as indicated (A) by the unpaired Student’s *t* test (A) or two-way ANOVA with the Bonferroni post hoc test (B and C).

**Figure S7.**
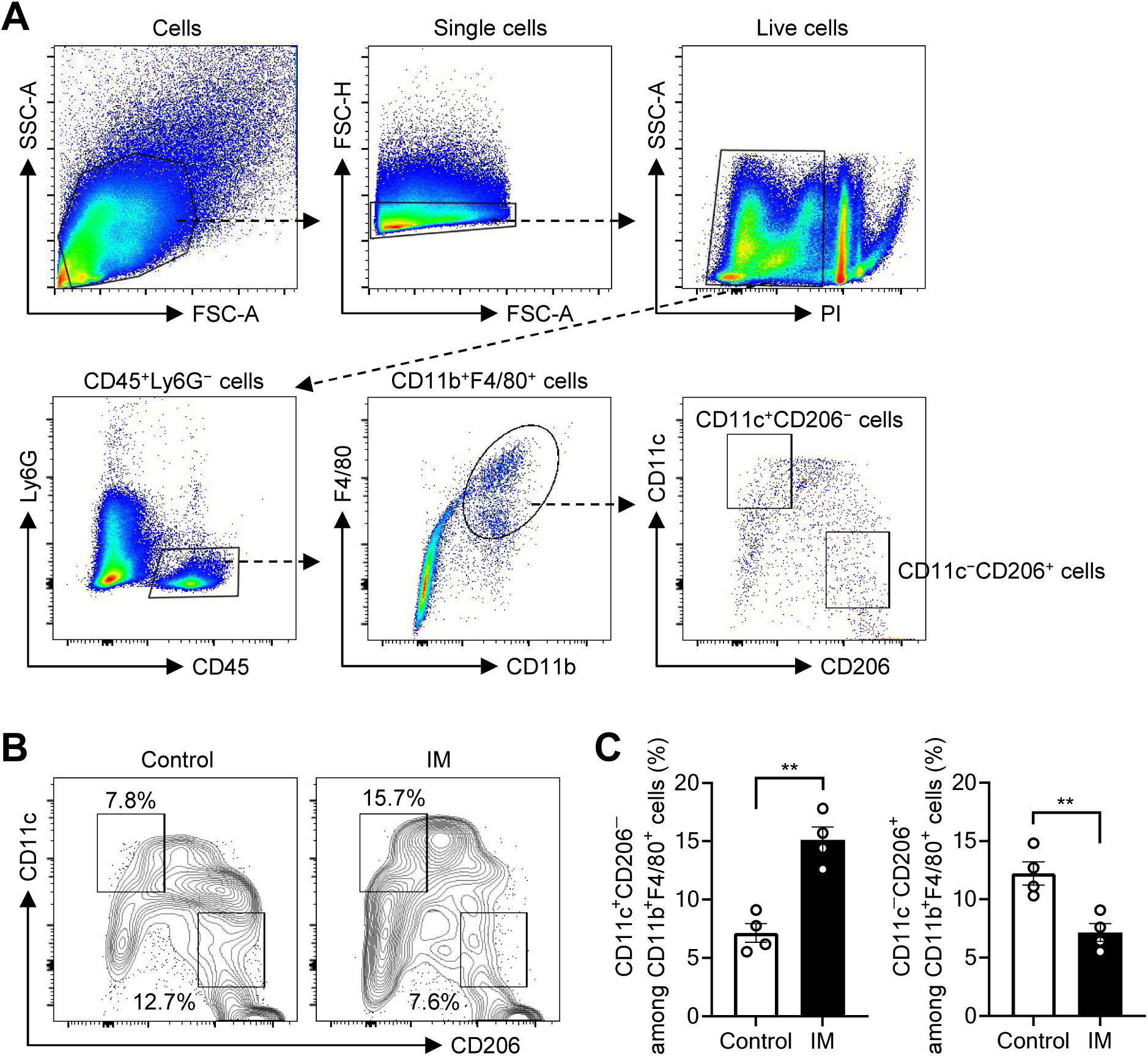
Gating strategy for macrophage subsets in the colon. (A) Gating strategy for fluorescence-activated sorting of macrophages from the colon of immobilized mice. (B and C) Representative flow cytometric analysis (B) and the percentage (C) of CD11c^+^CD206^−^ or CD11c^−^CD206^+^ cells in the colon of cast-immobilized (10 days) and control mice (*n* = 4 mice). Quantitative data are presented as means ± SEM (C). \*\**p* < 0.01 by the unpaired Student’s *t* test.

**Figure S8.**
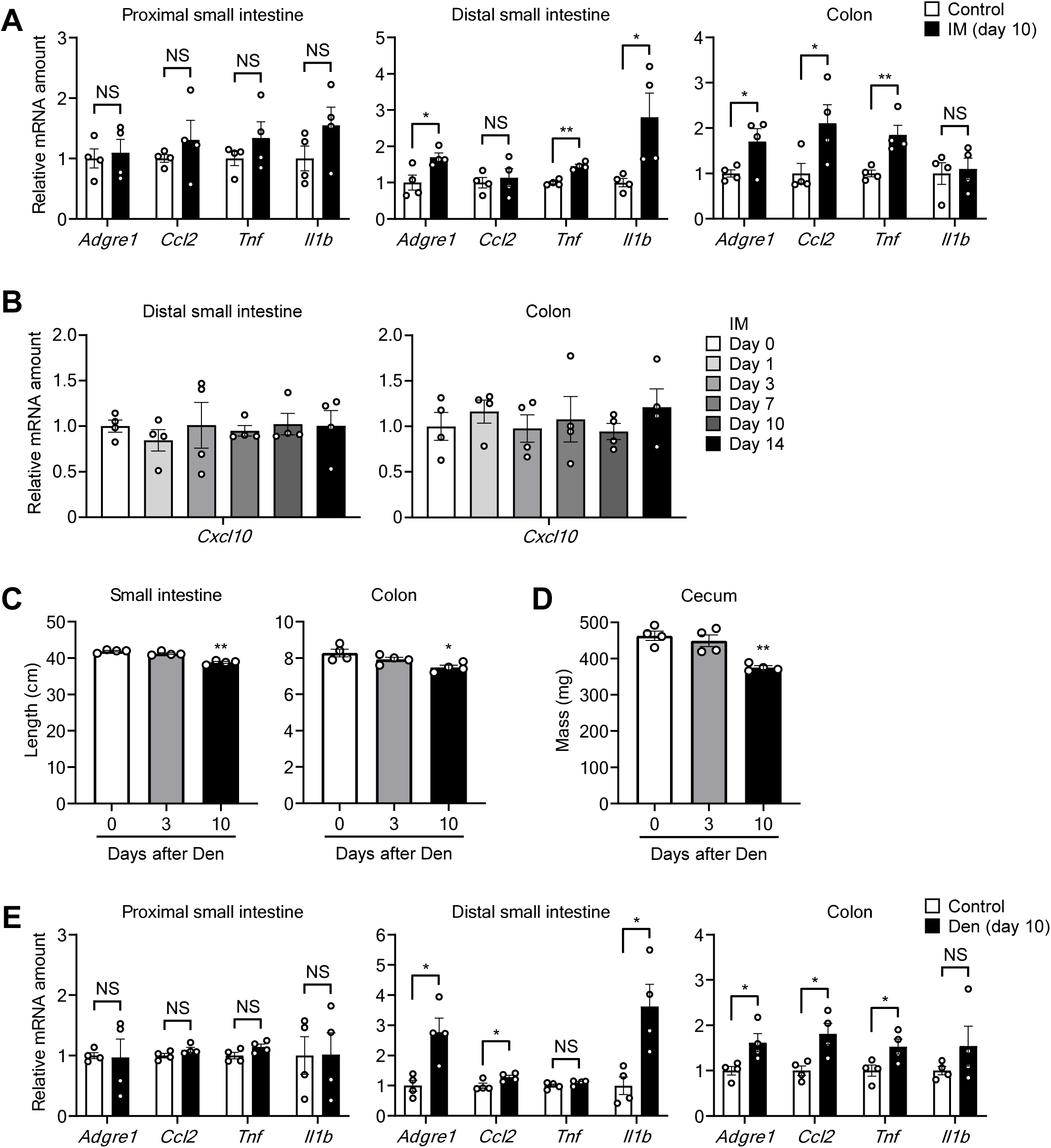
Effects of prolonged limb immobilization by cast-fixation or denervation on intestinal tissue. (A) RT-qPCR analysis of inflammation-related gene expression in the proximal or distal small intestine and colon of cast-immobilized (10 days) and control mice (*n* = 4 mice). (B) RT-qPCR analysis of *Cxcl10* mRNA abundance in the distal small intestine and colon of cast-immobilized mice (*n* = 4 mice). (C and D) Length of the small intestine and colon (C) as well as mass of the cecum (D) for mice subjected to limb denervation for the indicated times (*n* = 4 mice). (E) RT-qPCR analysis of inflammation-related gene expression in the proximal or distal small intestine and colon of denervated and control mice (*n* = 4 mice). Quantitative data are presented as means ± SEM (A–E). \**p* < 0.05, \*\**p* < 0.01, NS compared with day 0 (B–D) or as indicated (A and E) by the unpaired Student’s *t* test (A and E) or two-way ANOVA with the Bonferroni post hoc test (B–D).

**Figure S9.**
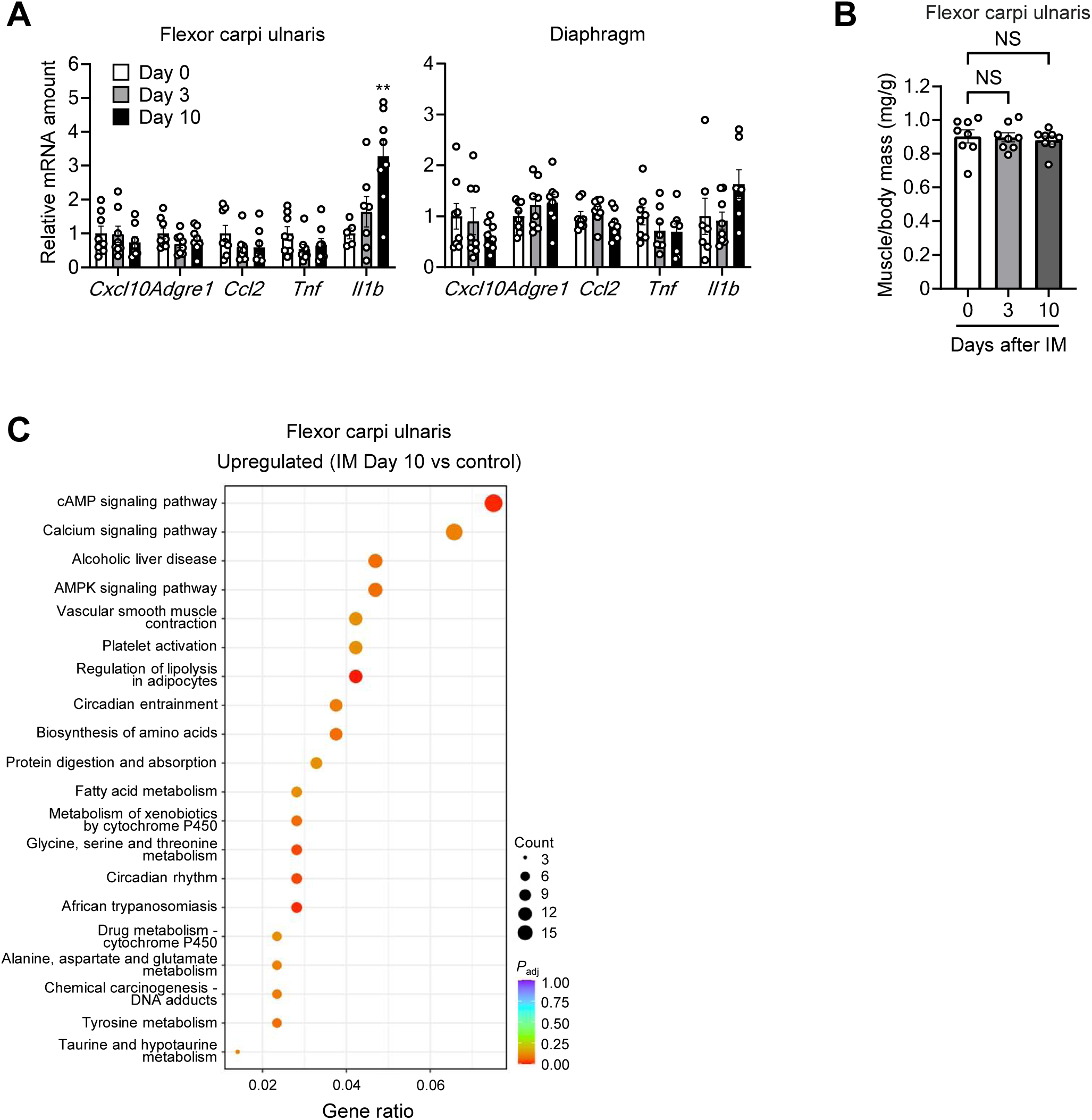
Time course of muscle mass and changes in gene expression in non-immobilized muscles. (A) RT-qPCR analysis of inflammation-related gene expression in flexor carpi ulnaris and diaphragm of immobilized mice (*n* = 8 mice). (B) Ratio of flexor carpi ulnaris mass to body mass for mice subjected to cast-immobilization for the indicated times (*n* = 8 mice). (C) KEGG pathway analysis for RNA-seq data from flexor carpi ulnaris at 10 days after hind limb immobilization. The top 20 pathways enriched among genes whose expression was upregulated by immobilization are shown. *p*_adj_, adjusted *p* value. Quantitative data are presented as means ± SEM (A and B). \*\**p* < 0.01, NS compared with day 0 (A) or as indicated (B) by two-way ANOVA with the Bonferroni post hoc test.

**Figure S10.**
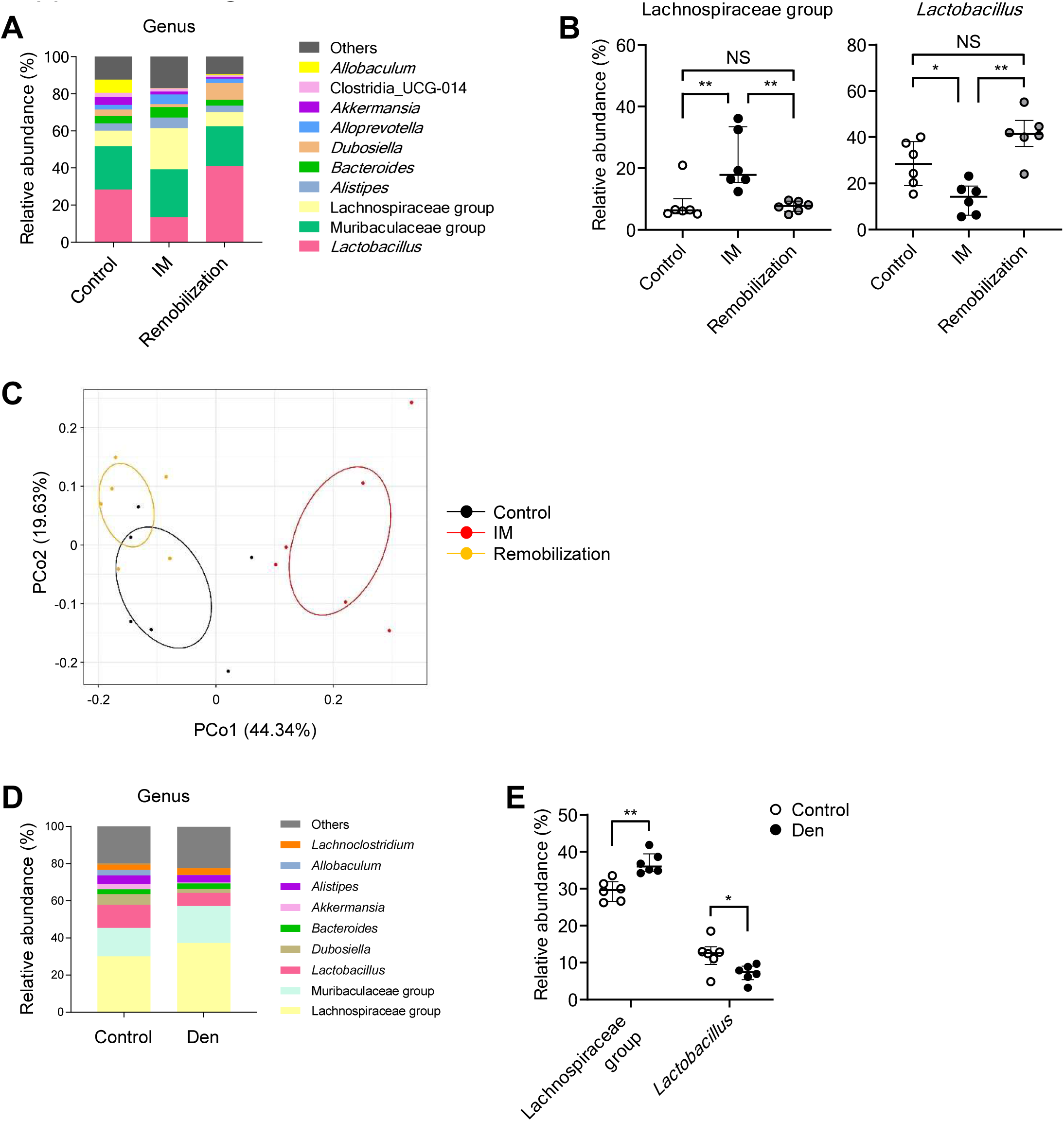
Effects of prolonged limb immobilization by cast-fixation or denervation on the gut microbiota. (A–C) Gut microbial composition at the genus level (A), relative abundance of the Lachnospiraceae group and *Lactobacillus* species (B), and corresponding principal coordinate analysis (C) for control mice and mice subjected to cast immobilization for 10 days with (remobilization) or without subsequent cast removal for 14 days (*n* = 6 mice). (D and E) Gut microbial composition at the genus level (D) and relative abundance of the Lachnospiraceae group and *Lactobacillus* species (E) for control mice or mice subjected to hind limb denervation for 10 days (*n* = 6 mice). Quantitative data are presented as medians with interquartile range (B and E). \**p* < 0.05, \*\**p* < 0.01, NS by the Mann-Whitney *U* test (E), one-way ANOVA with the Bonferroni post hoc test (B).

**Figure S11.**
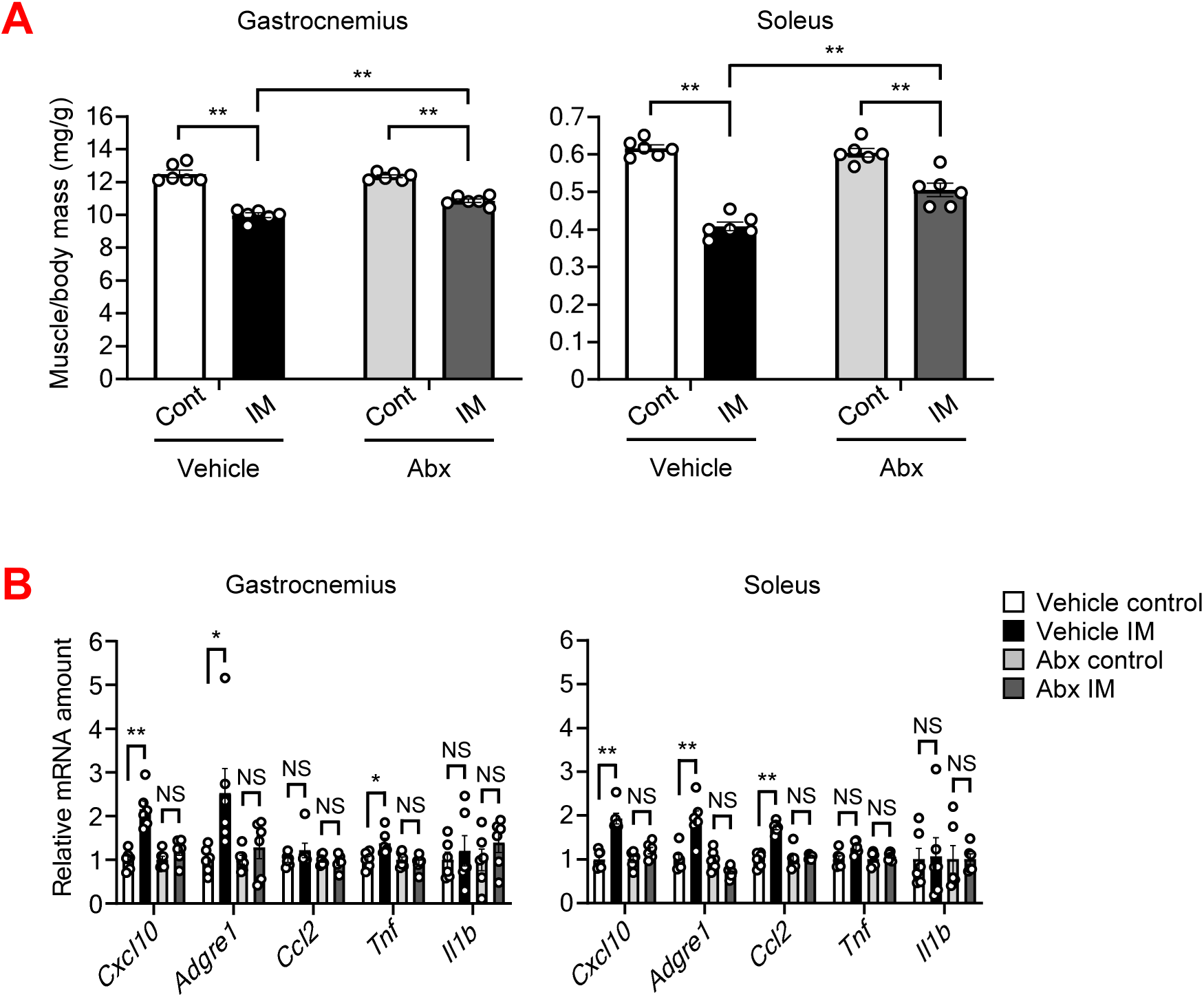
Effects of pseudosterilization of the intestine by antibiotics on muscle inflammation and atrophy induced by immobilization. (A and B) Ratio of gastrocnemius and soleus muscle mass to body mass (A), and RT-qPCR analysis of inflammation-related gene expression in gastrocnemius and soleus (B) of mice treated with a cocktail of antibiotics (Abx) or vehicle by oral gavage daily from 4 days before immobilization, with or without cast-immobilization for 10 days (*n* = 6 mice). Quantitative data are presented as means ± SEM. \**p* < 0.05, \*\**p* < 0.01, NS by two-way ANOVA with the Bonferroni post hoc test.

**Figure S12.**
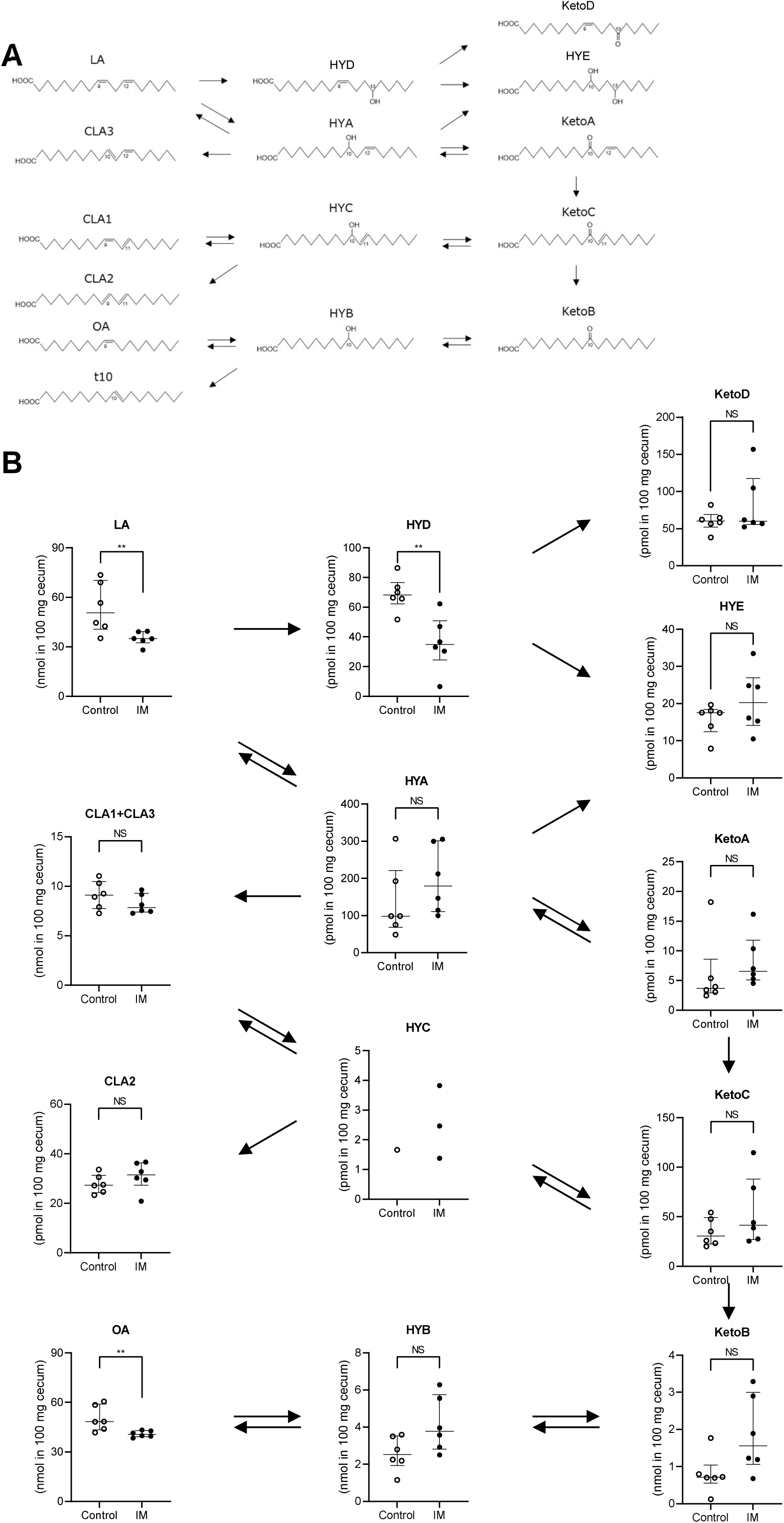
Prolonged limb immobilization by cast-fixation influences the fecal levels of fatty acid metabolites produced by the gut microbiota. (A) Metabolic pathway of LA showing the structure of fatty acid metabolites. (B) Fatty acid concentrations in cecal content of mice subjected (or not) to cast-immobilization for 10 days (*n* = 6 mice). HYC was below the detection limit in most samples. Quantitative data are presented as medians with interquartile range (B). \*\**p* < 0.01, NS by the Mann-Whitney *U* test.

**Figure S13.**
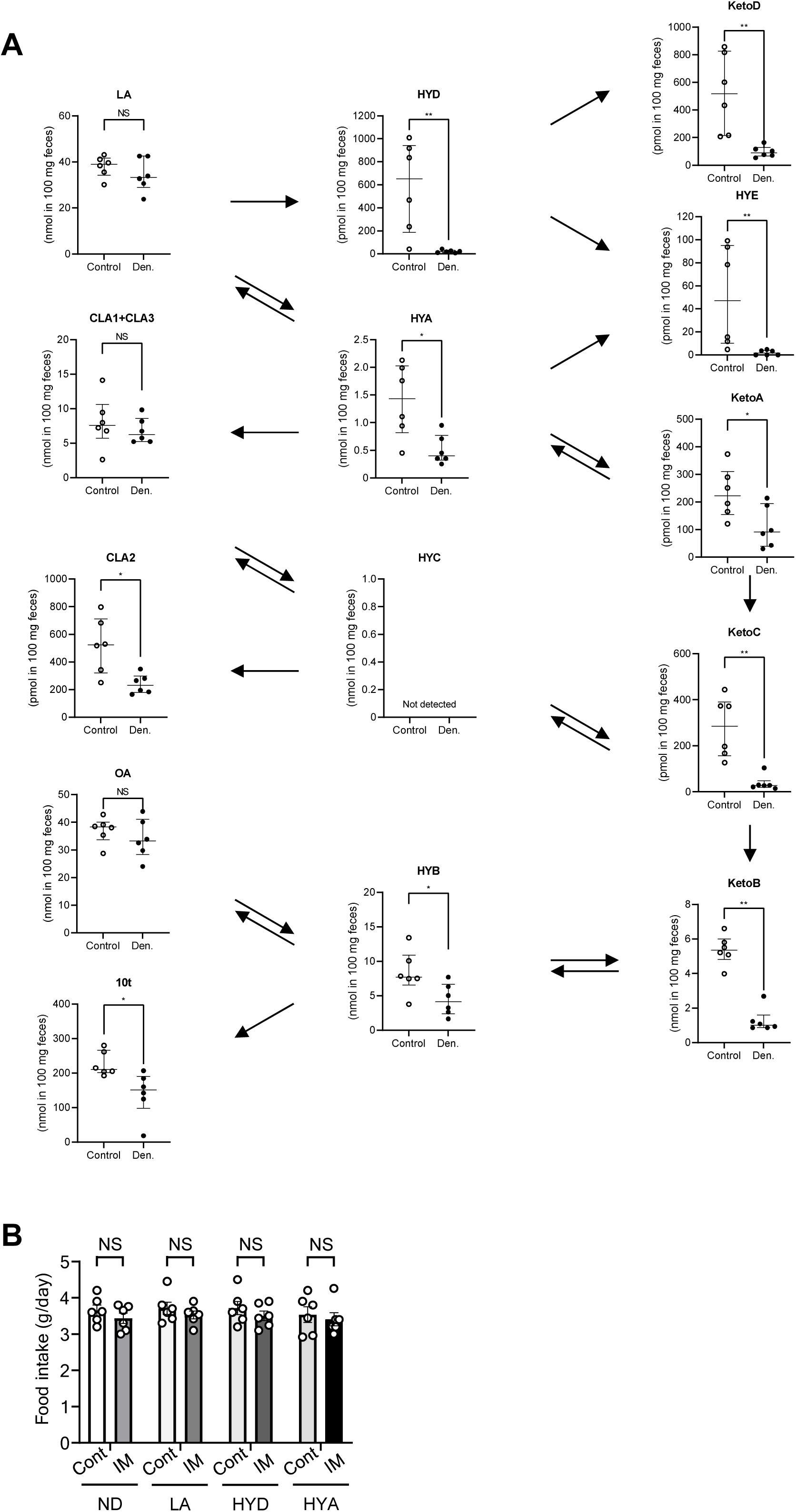
Prolonged limb immobilization by denervation influences the fecal levels of fatty acid metabolites produced by the gut microbiota. (A) Fatty acid concentrations in feces of control and denervated mice (10 days) (*n* = 6 mice). HYC was below the detection limit in all samples. (B) Daily food intake of mice maintained on a normal diet (ND) or a diet containing LA, HYD, or HYA and subjected (or not) to cast-immobilization for 10 days (*n* = 6 mice). Quantitative data are presented as means ± SEM (B) or medians with interquartile range (A). \**p* < 0.05, \*\**p* < 0.01, NS by the Mann-Whitney *U* test (A) or two-way ANOVA with the Bonferroni post hoc test (B).

**Figure S14.**
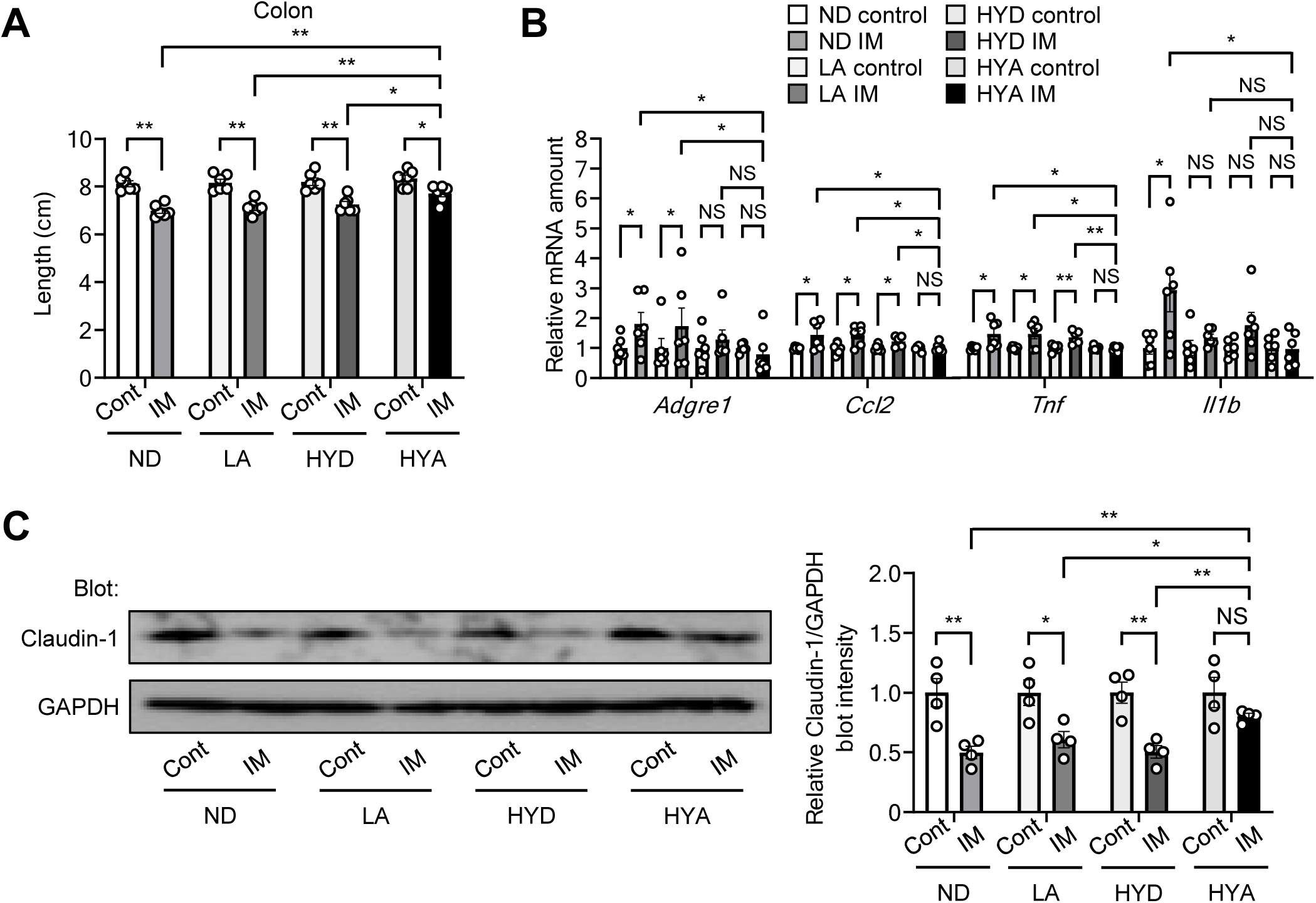
Effects of HYA on intestinal inflammation induced by prolonged limb immobilization. (A–C) Length of the colon (*n* = 6 mice) (A), RT-qPCR analysis of inflammation-related gene expression in the colon (*n* = 6 mice) (B), and immunoblot analysis of claudin-1 and GAPDH in the colon (*n* = 4 mice) (C) of mice fed a normal diet or a diet containing LA, HYD, or HYA and subjected (or not) to cast-immobilization for 10 days. Quantitative data are presented as means ± SEM (A–C). \**p* < 0.05, \*\**p* < 0.01, NS by two-way ANOVA with the Bonferroni post hoc test.

**Figure S15.**
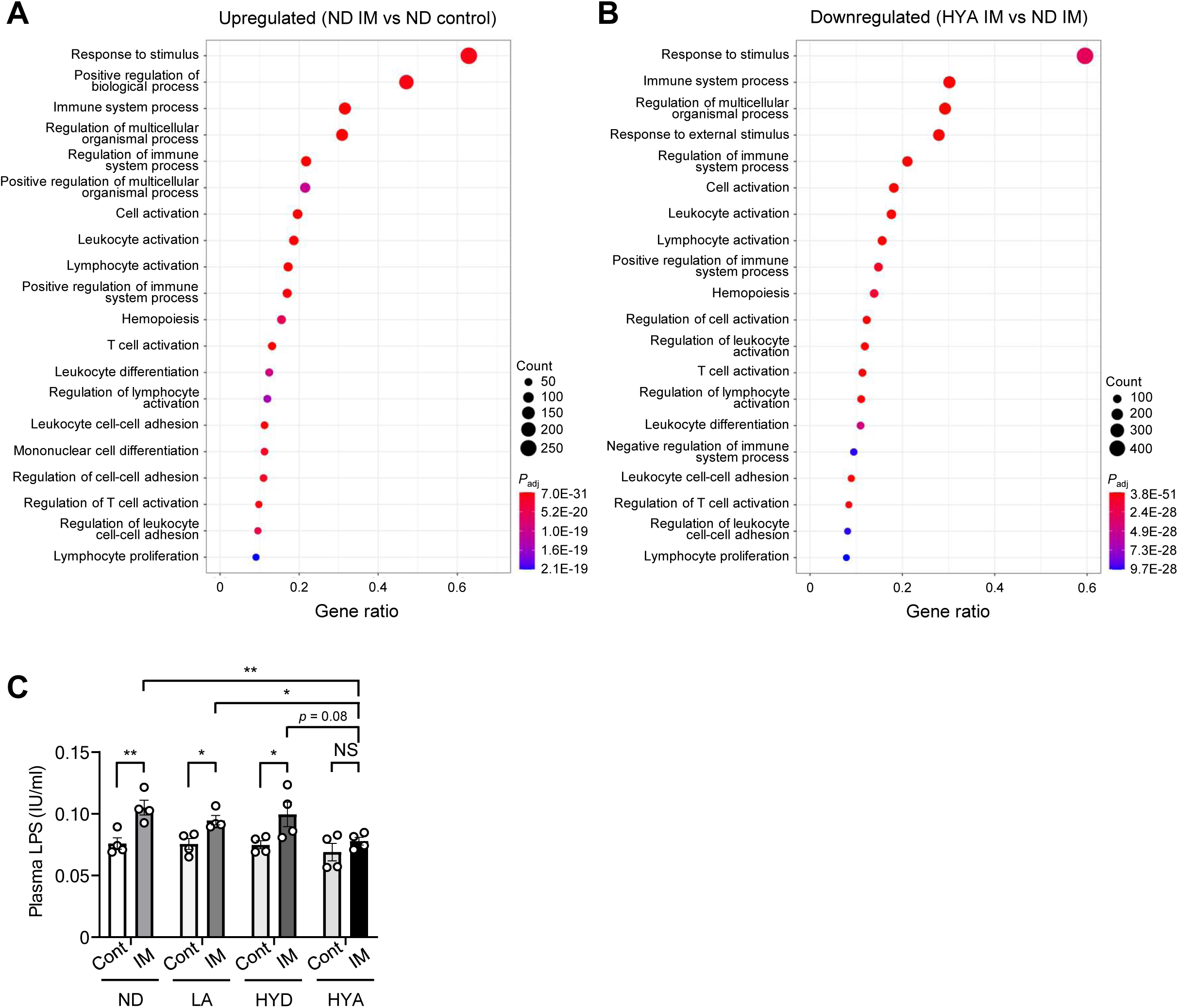
Effects of HYA on intestinal inflammation induced by prolonged limb immobilization. (A and B) GO functional analysis of RNA-seq data from the colon for biological processes upregulated in mice fed a normal diet and subjected to cast-immobilization for 10 days versus those not subjected to immobilization (A) as well as for biological processes downregulated in immobilized mice fed a diet containing HYA versus those fed a normal diet (B). The top 20 processes enriched among genes whose expression was altered by each condition are shown. *p*_adj_, adjusted *p* value. (C) Plasma concentration of LPS in mice fed a normal diet or a diet containing LA, HYD, or HYA and subjected (or not) to cast-immobilization for 10 days (*n* = 4 mice). Quantitative data are presented as means ± SEM (C). \**p* < 0.05, \*\**p* < 0.01, NS by two-way ANOVA with the Bonferroni post hoc test.

**Figure S16.**
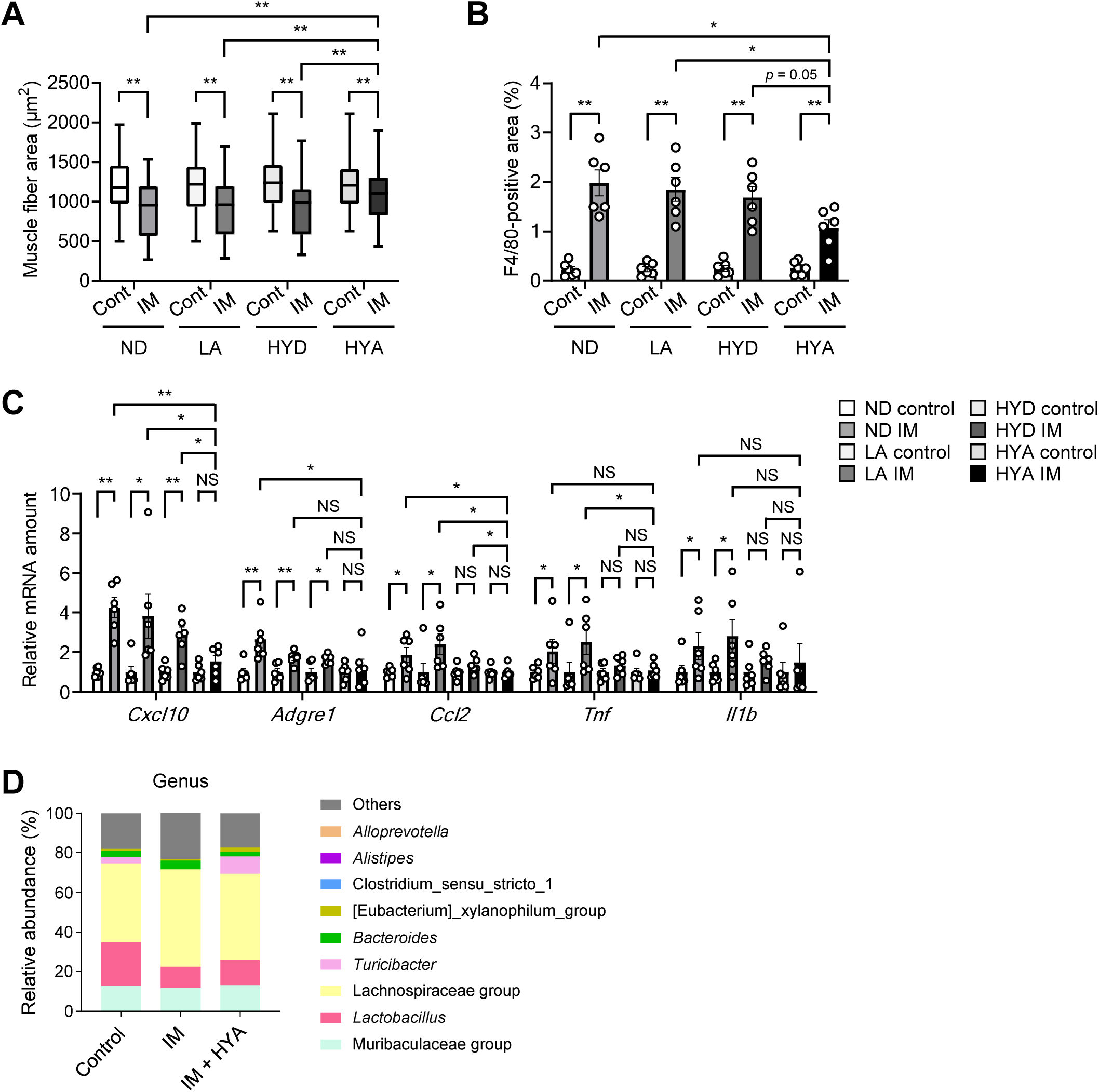
Effects of HYA on muscle inflammation and atrophy induced by prolonged limb immobilization. (A–C) Muscle fiber area (A) and the F4/80-positive area (B) in soleus (determined from images as in Figure 5E) as well as RT-qPCR analysis of inflammation-related gene expression in gastrocnemius (C) for mice fed a normal diet or a diet supplemented with LA, HYD, or HYA and subjected (or not) to cast-immobilization for 10 days (*n* = 6 mice). The area of 400 fibers pooled from four mice was measured and averaged for each condition in (A). (D) Gut microbial composition at the genus level for mice fed a normal diet or a diet containing HYA and subjected (or not) to cast-immobilization for 10 days (*n* = 4 mice). Quantitative data are presented as means ± SEM (B and C) or medians with interquartile range (A). \**p* < 0.05, \*\**p* < 0.01, NS by two-way ANOVA with the Bonferroni post hoc test.

**Figure S17.**
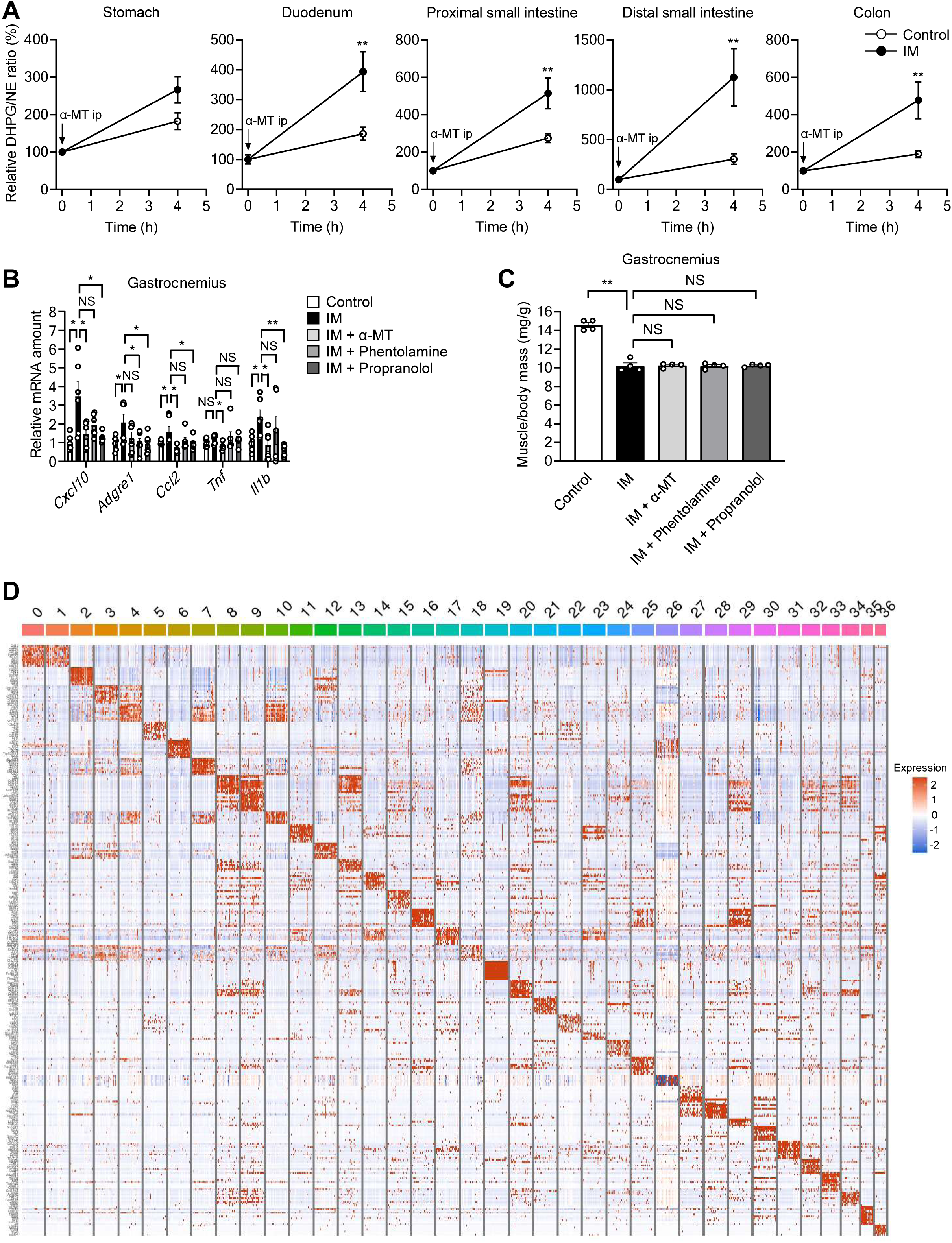
Sympathetic nerve activation contributes to muscle inflammation induced by limb immobilization. (A) Relative DHPG/NE ratio for the stomach, duodenum, proximal and distal small intestine, and colon of control or cast-immobilized (10 days) mice at 0 and 4 h after intraperitoneal (ip) injection of α-MT (*n* = 5 mice). (B and C) RT-qPCR analysis of inflammation-related gene expression in gastrocnemius (*n* = 6 mice) (B) as well as the ratio of gastrocnemius muscle mass to body mass (*n* = 4 mice) (C) for mice subjected to intraperitoneal injection of α-MT, phentolamine, propranolol, or vehicle daily for 10 days with or without concomitant cast-immobilization. (D) Heat map for the expression of all genes in the cell clusters of the colon identified by scRNA-seq analysis. Quantitative data are presented as means ± SEM (A–C). \**p* < 0.05, \*\**p* < 0.01, NS by two-way ANOVA with the Bonferroni post hoc test.

**Figure S18.**
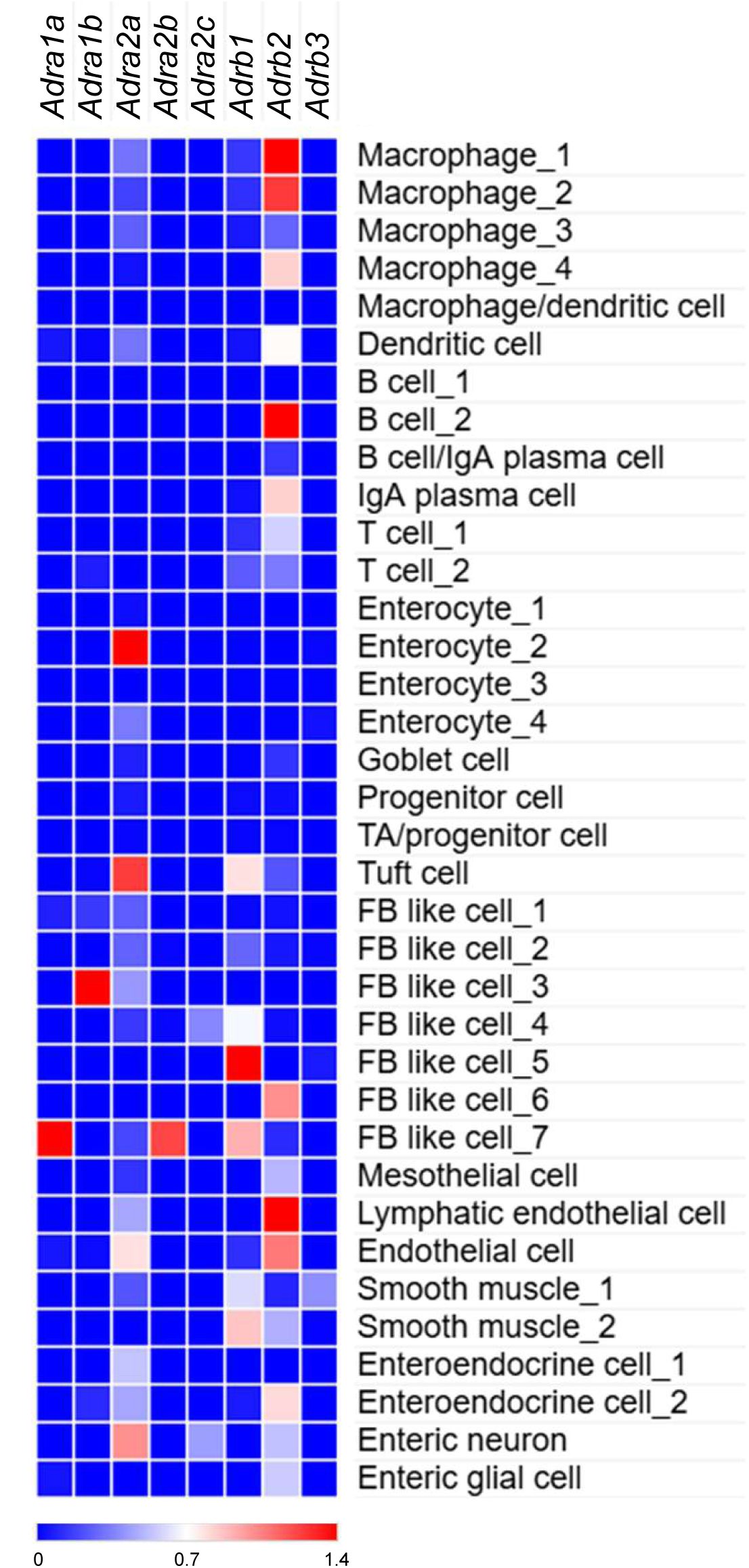
Heat map for expression of adrenergic receptor genes in cell clusters of the colon as determined by scRNA-seq analysis.

**Figure S19.**
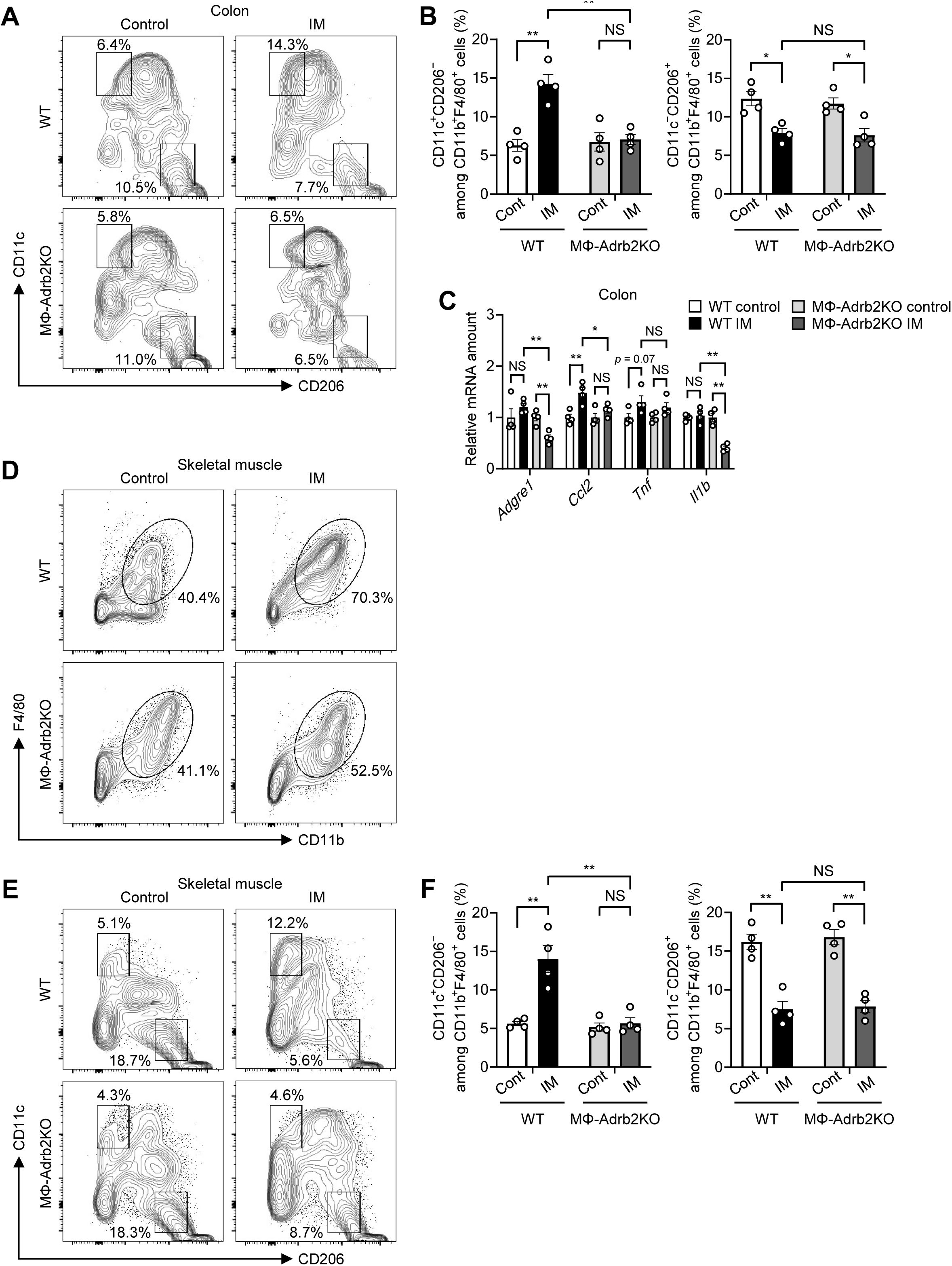
Intestinal and muscle inflammation induced by prolonged limb immobilization is attenuated in MΦ-Adrb2KO mice. (A–C) Representative flow cytometric analysis (A) and the percentage (B) of CD11c^+^CD206^−^or CD11c^−^CD206^+^ cells among CD11b^+^F4/80^+^ cells as well as RT-qPCR analysis of inflammation-related gene expression (C) in the colon of WT or MΦ-Adrb2KO mice subjected (or not) to cast-immobilization for 10 days (*n* = 4 mice). (D–F) Representative flow cytometric analysis of CD11b^+^F4/80^+^ cells among CD45^+^Ly6G^−^cells (D) as well as representative flow cytometric analysis (E) and the percentage (F) of CD11c^+^CD206^−^ or CD11c^−^CD206^+^ cells among CD11b^+^F4/80^+^ cells in hind limb skeletal muscle of WT or MΦ-Adrb2KO mice subjected (or not) to cast immobilization for 10 days (*n* = 4 mice). Quantitative data are presented as means ± SEM (B, C, and F). \**p* < 0.05, \*\**p* < 0.01, NS by two-way ANOVA with the Bonferroni post hoc test.

**Figure S20.**
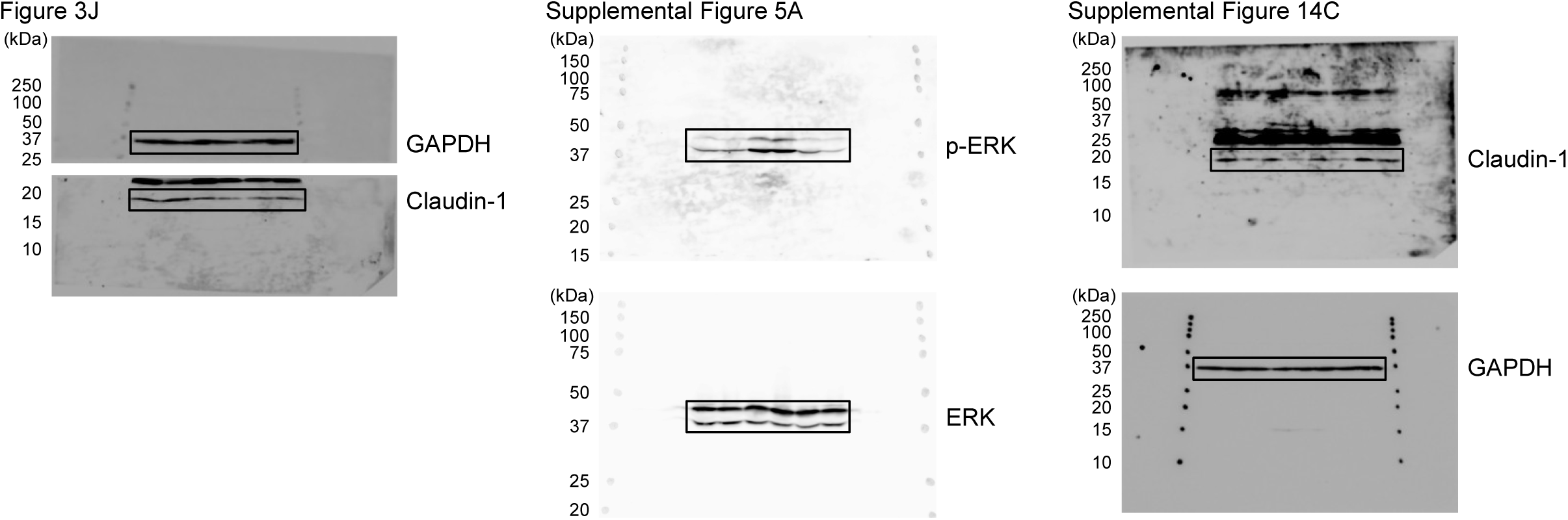
Uncropped immunoblot images. Black boxes contain the areas shown in the indicated figures.

**Table S1.**
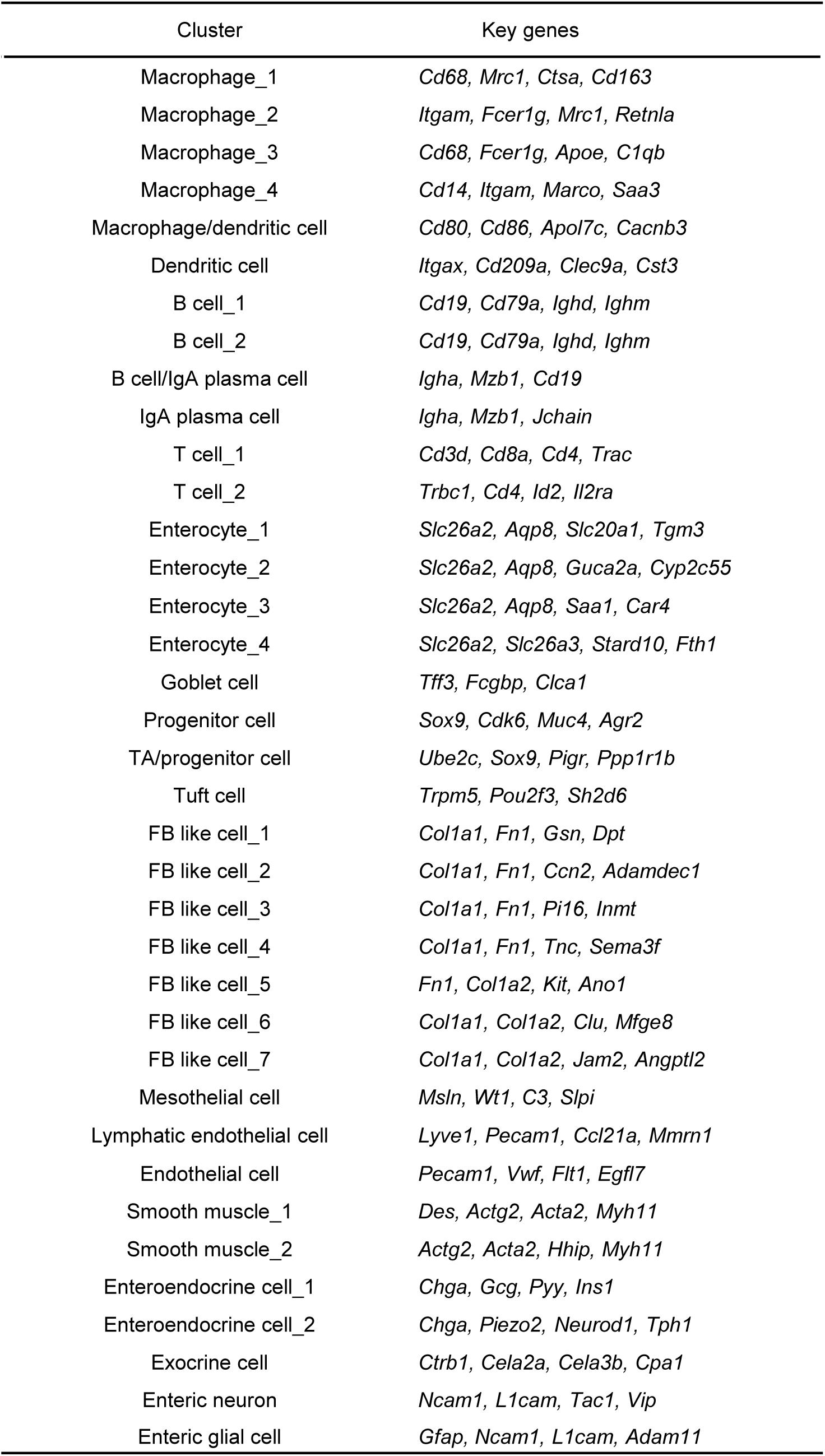
Genes that characterize cell clusters of the colon identified by scRNA-seq analysis of control mice and mice subjected to cast-immobilization for 10 days. The exocrine cell cluster is characterized by expression of genes for digestive enzymes including *Ctrb1*, *Cela2a*, *Cela3b*, and *Cap1*, all of which are commonly expressed in pancreatic exocrine cells. Moreover, this cluster was detected in a specific sample. Given that this cluster may have been derived from ectopic pancreatic tissue congenitally admixed with intestinal tissue, or from contaminating pancreatic tissue artefactually introduced during intestinal tissue collection, it was excluded from further analysis.

**Table S2.**
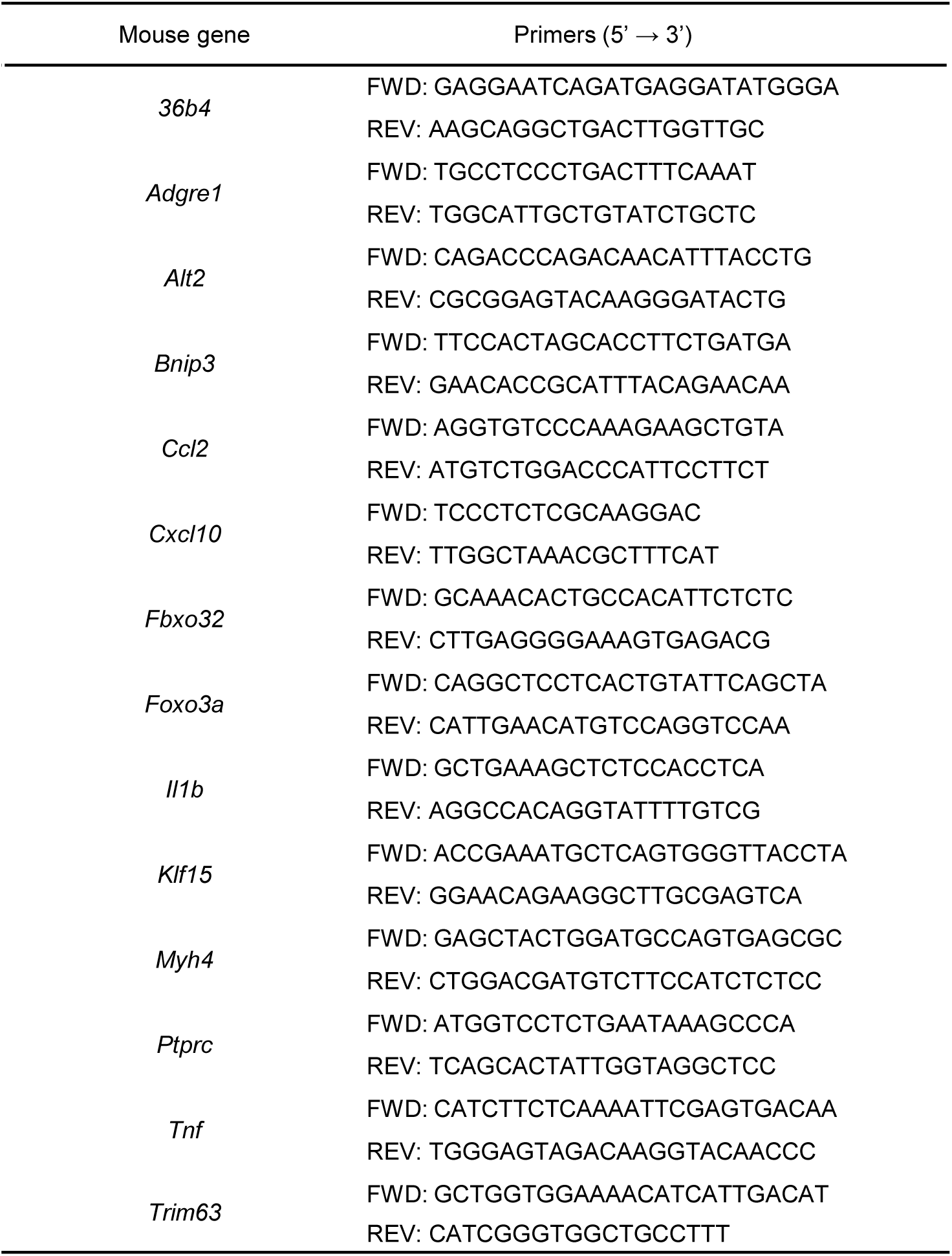
Sequences of mouse primers for RT-qPCR analysis. FWD, forward; REV, reverse.

**Table S3.**
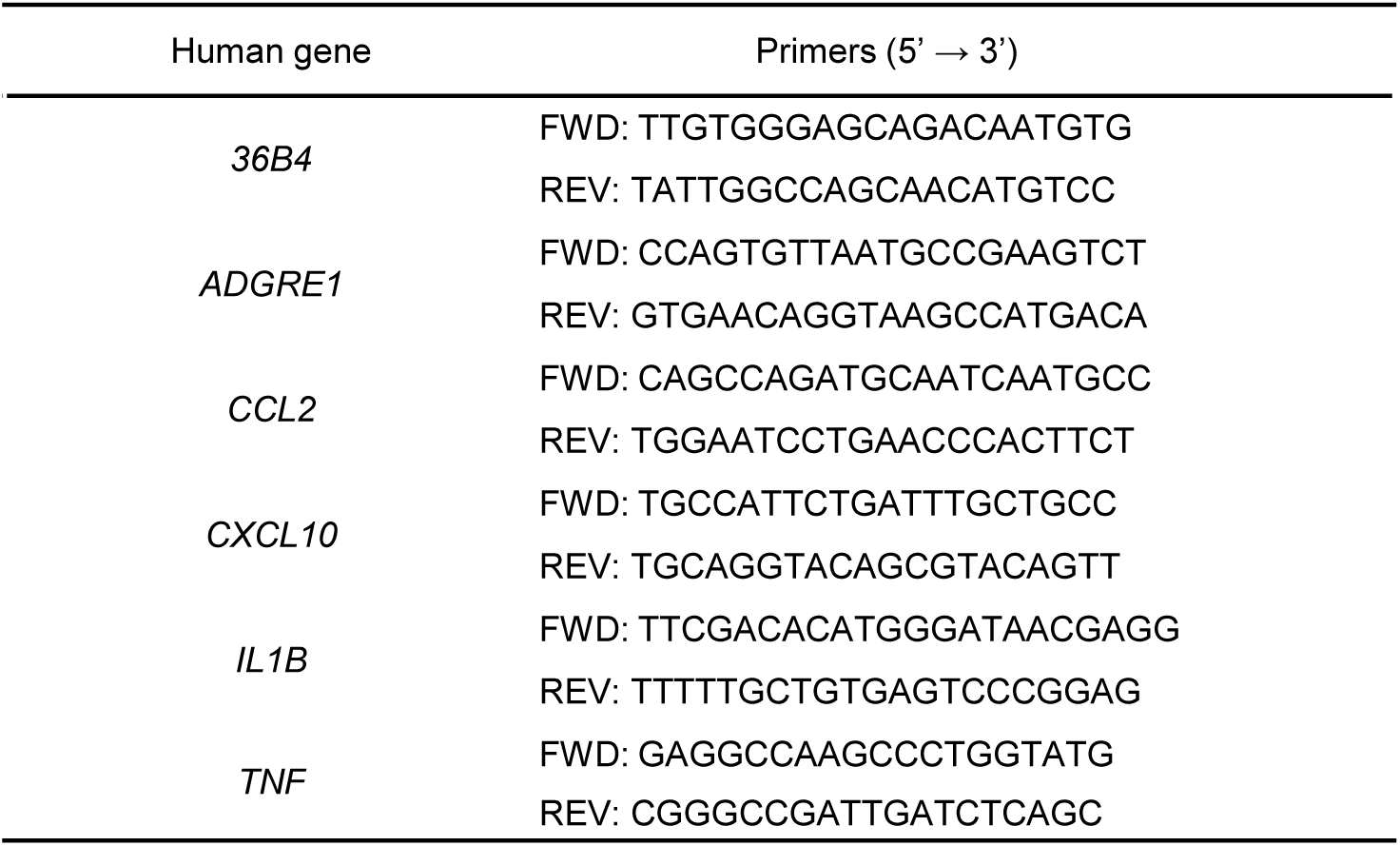
Sequences of human primers for RT-qPCR analysis. FWD, forward; REV, reverse.

## References

1. Kalyani, R.R., Corriere, M. & Ferrucci, L. Age-related and disease-related muscle loss: the effect of diabetes, obesity, and other diseases. Lancet Diabetes Endocrinol 2, 819–829 (2014).

2. Cartee, G.D., Hepple, R.T., Bamman, M.M. & Zierath, J.R. Exercise Promotes Healthy Aging of Skeletal Muscle. Cell Metab. 23, 1034–1047 (2016).

3. Preobrazenski, N., Seigel, J., Halliday, S., Janssen, I. & McGlory, C. Single-leg disuse decreases skeletal muscle strength, size, and power in uninjured adults: A systematic review and meta-analysis. J Cachexia Sarcopenia Muscle 14, 684–696 (2023).

4. Gao, Y., Arfat, Y., Wang, H. & Goswami, N. Muscle Atrophy Induced by Mechanical Unloading: Mechanisms and Potential Countermeasures. Front. Physiol. 9, 235 (2018).

5. Hardy, E.J.O., et al. The time course of disuse muscle atrophy of the lower limb in health and disease. J Cachexia Sarcopenia Muscle 13, 2616–2629 (2022).

6. Hirata, Y., et al. A Piezo1/KLF15/IL-6 axis mediates immobilization-induced muscle atrophy. J. Clin. Invest. 132, 1–13 (2022).

7. Saitou, K., et al. Local cyclical compression modulates macrophage function in situ and alleviates immobilization-induced muscle atrophy. Clin. Sci. (Lond.) 132, 2147–2161 (2018).

8. Kawanishi, N., Funakoshi, T. & Machida, S. Time-course study of macrophage infiltration and inflammation in cast immobilization-induced atrophied muscle of mice. Muscle Nerve 57, 1006–1013 (2018).

9. Hoff, P., et al. Effects of 60-day bed rest with and without exercise on cellular and humoral immunological parameters. Cell. Mol. Immunol. 12, 483–492 (2015).

10. Berchiche, Y.A. & Sakmar, T.P. CXC Chemokine Receptor 3 Alternative Splice Variants Selectively Activate Different Signaling Pathways. Mol. Pharmacol. 90, 483–495 (2016).

11. Kawano, Y., et al. Colonic Pro-inflammatory Macrophages Cause Insulin Resistance in an Intestinal Ccl2/Ccr2-Dependent Manner. Cell Metab. 24, 295–310 (2016).

12. Tilg, H., Zmora, N., Adolph, T.E. & Elinav, E. The intestinal microbiota fuelling metabolic inflammation. Nat. Rev. Immunol. 20, 40–54 (2020).

13. Garcia-Hernandez, V., Quiros, M. & Nusrat, A. Intestinal epithelial claudins: expression and regulation in homeostasis and inflammation. Ann. N. Y. Acad. Sci. 1397, 66–79 (2017).

14. Kishino, S., et al. Polyunsaturated fatty acid saturation by gut lactic acid bacteria affecting host lipid composition. Proc. Natl. Acad. Sci. U. S. A. 110, 17808–17813 (2013).

15. Miyamoto, J., et al. A gut microbial metabolite of linoleic acid, 10-hydroxy-cis-12-octadecenoic acid, ameliorates intestinal epithelial barrier impairment partially via GPR40-MEK-ERK pathway. J. Biol. Chem. 290, 2902–2918 (2015).

16. Hirata, A., et al. A novel unsaturated fatty acid hydratase toward C16 to C22 fatty acids from Lactobacillus acidophilus. J. Lipid Res. 56, 1340–1350 (2015).

17. Yang, D., Almanzar, N. & Chiu, I.M. The role of cellular and molecular neuroimmune crosstalk in gut immunity. Cell. Mol. Immunol. 20, 1259–1269 (2023).

18. Populin, L., Stebbing, M.J. & Furness, J.B. Neuronal regulation of the gut immune system and neuromodulation for treating inflammatory bowel disease. FASEB Bioadv 3, 953–966 (2021).

19. Saito, M., Minokoshi, Y. & Shimazu, T. Accelerated norepinephrine turnover in peripheral tissues after ventromedial hypothalamic stimulation in rats. Brain Res. 481, 298–303 (1989).

20. Shiuchi, T., et al. Hypothalamic orexin stimulates feeding-associated glucose utilization in skeletal muscle via sympathetic nervous system. Cell Metab. 10, 466–480 (2009).

21. Harima, Y., et al. Parallel labeled-line organization of sympathetic outflow for selective organ regulation in mice. Nat Commun 15, 10478 (2024).

22. Kuramoto, N., et al. Role of PDK1 in skeletal muscle hypertrophy induced by mechanical load. Sci. Rep. 11, 3447 (2021).

23. Lynch, G.S. & Ryall, J.G. Role of beta-adrenoceptor signaling in skeletal muscle: implications for muscle wasting and disease. Physiol. Rev. 88, 729–767 (2008).

24. Wang, X., Yu, L. & Wu, A.R. The effect of methanol fixation on single-cell RNA sequencing data. BMC Genomics 22, 420 (2021).

25. van Eijck, C.W.F., et al. Enhanced antitumour immunity following neoadjuvant chemoradiotherapy mediates a favourable prognosis in women with resected pancreatic cancer. Gut 73, 311–324 (2024).

26. Armet, A.M., et al. Rethinking healthy eating in light of the gut microbiome. Cell Host Microbe 30, 764–785 (2022).

27. Brown, E.M., Clardy, J. & Xavier, R.J. Gut microbiome lipid metabolism and its impact on host physiology. Cell Host Microbe 31, 173–186 (2023).

28. Miyamoto, J., et al. Gut microbiota confers host resistance to obesity by metabolizing dietary polyunsaturated fatty acids. Nat Commun 10, 4007 (2019).

29. Kaikiri, H., et al. Supplemental feeding of a gut microbial metabolite of linoleic acid, 10-hydroxy-cis-12-octadecenoic acid, alleviates spontaneous atopic dermatitis and modulates intestinal microbiota in NC/nga mice. Int. J. Food Sci. Nutr. 68, 941–951 (2017).

30. Kasahara, N., et al. A gut microbial metabolite of linoleic acid ameliorates liver fibrosis by inhibiting TGF-beta signaling in hepatic stellate cells. Sci. Rep. 13, 18983 (2023).

31. Yamada, M., et al. A bacterial metabolite ameliorates periodontal pathogen-induced gingival epithelial barrier disruption via GPR40 signaling. Sci. Rep. 8, 9008 (2018).

32. Moriyama, S., et al. beta(2)-adrenergic receptor-mediated negative regulation of group 2 innate lymphoid cell responses. Science 359, 1056–1061 (2018).

33. Wang, P., et al. Adrenergic nerves regulate intestinal regeneration through IL-22 signaling from type 3 innate lymphoid cells. Cell Stem Cell 30, 1166–1178 e1168 (2023).

34. Gabanyi, I., et al. Neuro-immune Interactions Drive Tissue Programming in Intestinal Macrophages. Cell 164, 378–391 (2016).

35. Matheis, F., et al. Adrenergic Signaling in Muscularis Macrophages Limits Infection-Induced Neuronal Loss. Cell 180, 64–78 e16 (2020).

36. Galzi, J.L., et al. Neutralizing endogenous chemokines with small molecules. Principles and potential therapeutic applications. Pharmacol. Ther. 126, 39–55 (2010).

37. van den Borne, P., Quax, P.H., Hoefer, I.E. & Pasterkamp, G. The multifaceted functions of CXCL10 in cardiovascular disease. Biomed Res Int 2014, 893106 (2014).

38. Sasaki, Y., et al. Blockade of the CXCR3/CXCL10 axis ameliorates inflammation caused by immunoproteasome dysfunction. JCI Insight 7(2022).

39. Yellin, M., et al. A phase II, randomized, double-blind, placebo-controlled study evaluating the efficacy and safety of MDX-1100, a fully human anti-CXCL10 monoclonal antibody, in combination with methotrexate in patients with rheumatoid arthritis. Arthritis Rheum. 64, 1730–1739 (2012).

40. Alex Klarenbeek, D.M., Christophe Blanchetot, Michael Saunders, Sebastian van der Woning, Martine Smit, Hans de Haard, Erik Hofman. Targeting chemokines and chemokine receptors with antibodies. Drug Discov Today Technol 9, e227–314 (2012).

41. Hinoi, E., et al. The sympathetic tone mediates leptin’s inhibition of insulin secretion by modulating osteocalcin bioactivity. J. Cell Biol. 183, 1235–1242 (2008).

42. Clausen, B.E., Burkhardt, C., Reith, W., Renkawitz, R. & Forster, I. Conditional gene targeting in macrophages and granulocytes using LysMcre mice. Transgenic Res. 8, 265–277 (1999).

43. Wouters, S., et al. Protocol for fecal microbiota transplantation: A microaerophilic approach for mice housed in a specific pathogen-free facility. STAR Protoc 6, 103517 (2025).

44. Rohm, T.V., et al. Targeting colonic macrophages improves glycemic control in high-fat diet-induced obesity. Commun Biol 5, 370 (2022).

45. Tyus, D., et al. The sympathetic nervous system drives hyperinflammatory responses to Clostridioides difficile infection. Cell Rep Med 5, 101771 (2024).

46. Hirata, Y., et al. Hyperglycemia induces skeletal muscle atrophy via a WWP1/KLF15 axis. JCI Insight 4(2019).

47. Prieto, C. & Barrios, D. RaNA-Seq: Interactive RNA-Seq analysis from FASTQ files to functional analysis. Bioinformatics (2019).

48. Tsuji, K., Shimada, W., Kishino, S., Ogawa, J. & Arita, M. Comprehensive analysis of fatty acid metabolites produced by gut microbiota using LC-MS/MS-based lipidomics. Medical Mass Spectrometry 6, 112–125 (2022).

